# Haplotype-based analysis distinguishes maternal-fetal genetic contribution to pregnancy-related outcomes

**DOI:** 10.1101/2020.05.12.079863

**Authors:** Amit K. Srivastava, Julius Juodakis, Pol Sole-Navais, Jing Chen, Jonas Bacelis, Kari Teramo, Mikko Hallman, Pal R. Njølstad, David M. Evans, Bo Jacobsson, Louis J. Muglia, Ge Zhang

## Abstract

Genotype-based approaches for the estimation of SNP-based narrow-sense heritability 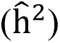 have limited utility in pregnancy-related outcomes due to confounding by the shared alleles between mother and child. Here, we propose a haplotype-based approach to estimate the genetic variance attributable to three haplotypes – maternal transmitted 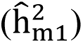, maternal non-transmitted 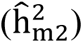 and paternal transmitted 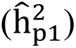 in mother-child pairs. We show through extensive simulations that our haplotype-based approach outperforms the conventional and contemporary approaches for resolving the contribution of maternal and fetal effects, particularly when m1 and p1 have different effects in the offspring. We apply this approach to estimate the explicit and relative maternal-fetal genetic contribution to the phenotypic variance of gestational duration and gestational duration adjusted fetal size measurements at birth in 10,375 mother-child pairs. The results reveal that variance of gestational duration is mainly attributable to m1 and m2 (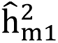= 17.3%, S. E. = 5.2%; 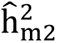 = 12.2%, S. E. = 5.2%; 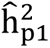 = 0.0%, S. E. = 5.0%). In contrast, variance of fetal size measurements at birth are mainly attributable to m1 and p1 (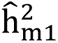 = 18.6 − 36.4%, 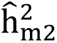 = 0.0 − 5.2% and 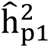 = 4.4 − 13.6%). Our results suggest that gestational duration and fetal size measurements are primarily genetically determined by the maternal and fetal genomes, respectively. In addition, a greater contribution of m1 as compared to m2 and p1 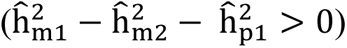 to birth length and head circumference suggests a substantial influence of correlated maternal-fetal genetic effects on these traits. Our newly developed approach provides a direct and robust alternative for resolving explicit maternal and fetal genetic contributions to the phenotypic variance of pregnancy-related outcomes.

## Introduction

Narrow sense heritability (h^2^) is the proportion of phenotypic variance in a population attributable to additive genetic values (breeding values)^1^. Generally, the concept of the h^2^ estimation comes from balanced designs – regression of a child’s phenotype on mid-parent phenotype, correlation of full or half sibs and differences in the correlation of monozygotic and dizygotic twins^1^. However, in a population with mixed relationships, linear mixed model (LMM) is the most flexible approach accounting for both fixed and random effects^1–5^.

Over the last decade, various methods^6^ including Genome-based Restricted Maximum Likelihood (GREML)^7,8^, Linkage Disequilibrium Adjusted kinships (LDAK)^9^, threshold Genomic Relatedness Matrices (Threshold-GRMs)^10^, LD Score regression (LDSC)^11^ and Phenotype Correlation-Genotype Correlation (PCGC)^12^ have been developed to estimate SNP-based narrow-sense heritability (commonly known as SNP-based heritability or SNP-heritability – 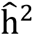)^13^. In addition, variants of these approaches such as GREML-MAF stratified (GREML-MS)^14^, GREML-LD and MAF stratified (GREML-LDMS)^15^ and LDAK-MAF stratified (LDAK-MS)^16^ have enabled partitioning of the genetic variance into additive and non-additive components as well as variance components attributable to chromosomes, genes and inter-genic regions. The above approaches have helped explain a large proportion of the missing heritability in various complex diseases and quantitative traits^8–11,13,16–18^. Nevertheless, conventional approaches utilizing an individual’s genotype information are less suited for pregnancy-related outcomes which are jointly influenced by direct fetal and indirect parental genetic effects^19–22^. In recent years, several studies using genotype information in mother-child duos ^19,21,23–26^ and parent-child trios^27^ have examined the contribution of parental genetic effects^28,29^ and fetal genetic effects in various pregnancy-related outcomes. However, these approaches are based on several assumptions including equal effects of maternal and paternal transmitted alleles in child. Hence, the estimation of heritability in pregnancy-related outcomes demand a direct approach with relaxed assumptions.

Here, we consider mother-child pair as a single analytical unit consisting of three haplotypes corresponding to maternal transmitted (m1), maternal non-transmitted (m2) and paternal transmitted (p1) alleles^30–32^. Use of such an analytical unit provides an advantage over conventional approaches based on individual’s genotype information by avoiding the confounding of m1 which can influence pregnancy-related outcomes through both the mother and child (Figure 1a)^22^. We generate three separate genetic relatedness matrices M1, M2 and P1 using only m1, only m2 and only p1, respectively. We fit all three matrices simultaneously in a linear mixed model (LMM) to estimate variance attributable to each haplotype (Figure 1b). Although our approach doesn’t directly estimate SNP-heritability, we use 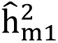, 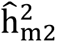 and 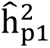 to represent variance attributable to m1, m2 and p1 respectively for the comparison purposes. We compare the behavior of our newly developed haplotype-based genome-wide complex trait analysis approach (H-GCTA) with existing genotype-based approaches such as Genome-wide Complex Traits Analysis (GCTA) and Maternal-Genome-wide Complex Traits Analysis (M-GCTA) approach using simulated phenotypes with varying contributions and correlations of maternal and fetal genetic effects. We show that H-GCTA outperforms the conventional and other contemporary approaches, particularly when the maternal and paternal transmitted alleles have different effects (e.g. parent-of-origin effects – POEs) on a fetal trait.

**Figure 1:**
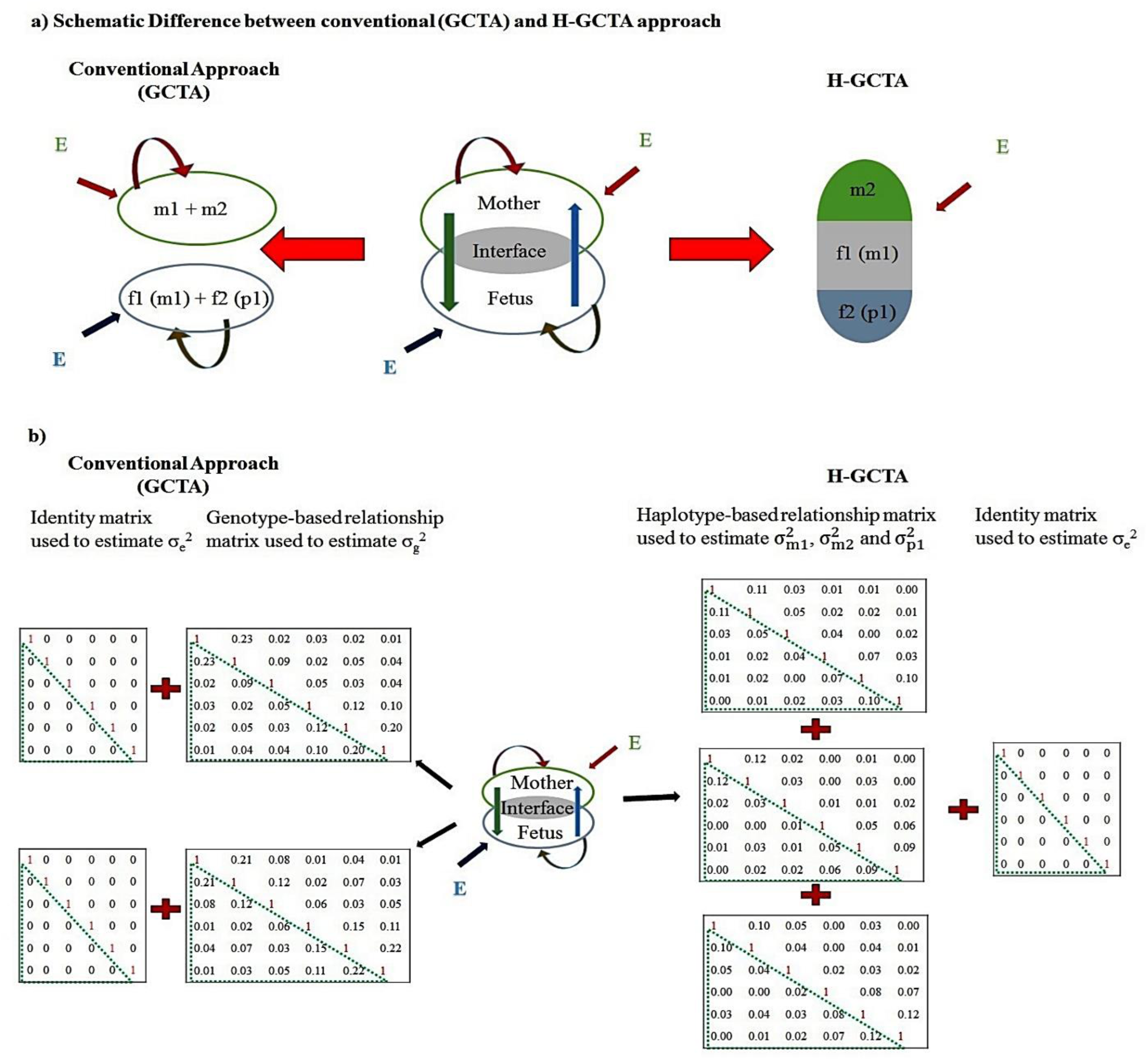
a) Schematic representation of the difference between the conventional genotype-based and newly developed haplotype-based analysis approach; the left part of the figure represents the conventional approach based on genotypes of mother and child separately and the right part represents haplotype-based analysis by treating mother/child pairs as analytical units. Green vertical arrow represents maternal genetic effects in fetus, whereas Blue one represents fetal genetic effects in mother during pregnancy. Red and Golden curved arrows represent maternal and fetal genetic effects in mother and fetus, respectively. Red and Indigo slant arrows represent the environmental effects on mother and fetus, respectively. m1 (f1): Maternal transmitted alleles; m2: Maternal non-transmitted alleles; f2 (p1): paternal transmitted alleles; E: Environmental factors. b) Schematic representation of the difference between conventional approach of heritability estimation utilizing genotype-based GRMs and our approach utilizing haplotype-based GRMs (representing the example of mother-child duos). While Conventional GCTA approach fits individual’s genotype-based GRM separately in mothers and children (left side), haplotype-based approach fits three haplotype-based GRMs together (right side). 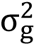: phenotypic variance attributable to mothers’ or children’s genotypes; 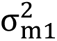, 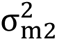 and 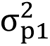: phenotypic variance attributable to m1, m2 and p1 respectively; 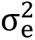: phenotypic variance attributable to E.

We further apply our approach to a cohort of 10,375 mother-child pairs to estimate the explicit and relative contribution of maternal-fetal genetic effects to the phenotypic variance of gestational duration and gestational duration adjusted fetal size measurements at birth, including birth weight, birth length and head circumference. Our results suggest that genetic variance in gestational duration is primarily attributable to the maternal genome, i.e. the maternal transmitted (m1) and non-transmitted (m2) alleles, whereas genetic variance in fetal size measurements at birth are largely attributable to fetal genome, i.e. maternal transmitted (m1) and paternal transmitted (p1) alleles. In addition, a higher attribution to m1 as compared to m2 and p1 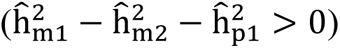 suggests a large contribution of correlated maternal-fetal genetic effects to the variance of birth length and head circumference. Our haplotype-based approach provides a direct method with relaxed underlying assumptions to estimate the explicit and relative maternal-fetal contributions to the phenotypic variance of pregnancy-related outcomes.

## Results

### Heritability Estimation using simulated data

We first evaluated the utility and robustness of H-GCTA using simulated phenotypes based on the real genotype data from a homogenous cohort (Avon Longitudinal Study of Parents And Children; ALSPAC) with 5,369 mother-child pairs and pooled dataset (diverse European populations, including ALSPAC) with 10,375 mother-child pairs. Traits were simulated with varying contributions and correlation of maternal and fetal genetic effects (Table 1 and methods). All traits were simulated with a total genetic variance at 50%, using a randomly selected set of 10,000 causal variants. Traits with correlated maternal-fetal genetic effects were simulated using the same set of causal variants in mother and child. In addition, we also incorporated different levels of POEs (maternal and paternal transmitted alleles had different effects in fetus) in varying proportion of causal variants for traits with only fetal effects (Table 1 and methods).

**Table 1:** Genetic models for simulation conditions – simulated maternal traits, fetal traits, traits with joint maternal-fetal effects and fetal traits with different levels of parent-of-origin effects (POEs), incorporated in varying proportion of causal variants. Correlated maternal-fetal effects were randomly drawn from multivariate normal distribution 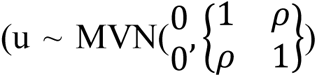 where, ρ represents correlation of maternal and fetal genetic effects which is equal to covariance of maternal-fetal genetic effects in a standard normal distribution. POEs were incorporated by reducing the effect of maternal transmitted alleles (m1) in comparison to paternal transmitted alleles (p1) by multiplying effects of m1 with (1 – I) where I is the imprinting factor such as 0.25, 0.50, 0.75 and 1.0. In case of I =1.0, m1 has no effect which represents complete imprinting whereas other I values represent partial imprinting. u represents the allelic effects; g_m1_ and g_m2_ represent the vectors corresponding to maternal transmitted and non-transmitted alleles in mother whereas g_f1_ and g_f2_ represent the vectors corresponding to maternal and paternal transmitted alleles in child, respectively. It is noteworthy that g_m1_ and g_f1_ are same in phased mother-child pairs. Likewise, f2 is represented as p1 (paternal transmitted alleles) throughout the article. A total of 100 randomly picked residual values 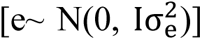 were added to genetic values to generate 100 replicates of each simulated phenotype (see methods).

| Causal effects | Genetic values | Notes |
| --- | --- | --- |
| Only maternal | $(g_{m1} * u_{m1}) + (g_{m2} * u_{m2})$ | $u_{m1} = u_{m2} = u_m$ |
| Only fetal | $(g_{f1} * u_{f1}) + (g_{f2} * u_{f2})$ | $u_{f1} = u_{f2} = u_f$ |
| Joint maternal-fetal | $(g_{m1} * u_{m1}) + (g_{m2} * u_{m2}) + (g_{f1} * u_{f1}) + (g_{f2} * u_{f2})$ | $u_{m1} = u_{m2} = u_m; u_{f1} = u_{f2} = u_f$ with average $\text{cor}(u_m, u_f) = \rho$, where $\rho = \{0.0, -0.5, -1.0, 0.5, 1.0\}$. For $\rho = 0.0$, same set and independent sets of causal variants were selected from mothers and fetuses. |
| Only fetal with POEs | $(g_{f1} * u_{f1}) + (g_{f2} * u_{f2})$ | $u_{f1} = (1 - I) * u_{f2}$, where $I = \{0.25, 0.50, 0.75, 1.0\}$. POEs were introduced to different proportions of causal variants. |

We compared the performance of H-GCTA with conventional GCTA approach and a contemporary M-GCTA approach for each simulated trait. For each approach, we estimated the genetic variance using three models – GREML, LDAK-Thin (where all pruned SNPs were given equal weights) and LDAK with SNP-specific weights (hereafter referred as LDAK-Weights, where each SNP had different weights based on pair-wise LD). Using any particular approach, each model yielded similar results when used with recommended α values (GREML: α = –1.0; LDAK and LDAK-Thin: α = –0.25) which represents the extent to which minor allele frequency (MAF) influences the variance of SNP effects on phenotypes^16^. We observed that the estimated genetic variance were similar in pooled datasets and homogenous cohort ALSPAC. However, due to small sample size, the estimated genetic variance in ALSPAC cohort had larger standard errors (Supplementary Figures 3, 4 and Supplementary Tables 5-12). Here, we discuss the results of simulated traits from pooled dataset using GREML (α = –1.0) fitted through GCTA, M-GCTA and H-GCTA approach.

### Heritability of simulated traits with only maternal effects

Using conventional approach for maternal traits in mothers and children separately, the estimated SNP-heritability (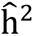) based on maternal (m) and fetal (f) genotypes was 45.0% (S.E. = 8.6%) and 13.6% (S.E. = 8.6%) respectively (Figure 2a, Supplementary Table 13). We also used M-GCTA in mother-child duos to estimate the variance attributable to indirect maternal effect (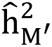 = 40.6%, S. E. = 6.1%), direct fetal effect (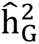 = −2.2%, S. E. = 6.0%) and maternal-fetal covariance (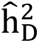 = 5.4%, S. E. = 5.2%) (Figure 2a, Supplementary Table 13). Using H-GCTA for maternal traits in mother-child duos, variance attributable to maternal transmitted alleles (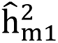), maternal non-transmitted alleles (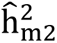), and paternal transmitted alleles (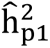) were 25.5% (S.E. = 4.8%), 22.9% (S.E. = 4.5%) and –3.0% (S.E. = 4.2%) respectively (Figure 2a, Supplementary Table 13). M-GCTA and H-GCTA accurately distinguished the maternal origin of the simulated traits; however, the conventional GCTA also showed a superficial contribution from the fetal genome (13.6%, approximately one quarter of the 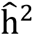 based on maternal genotype) due to 50% alleles shared between mother and child (Supplementary Table 13).

**Figure 2:**
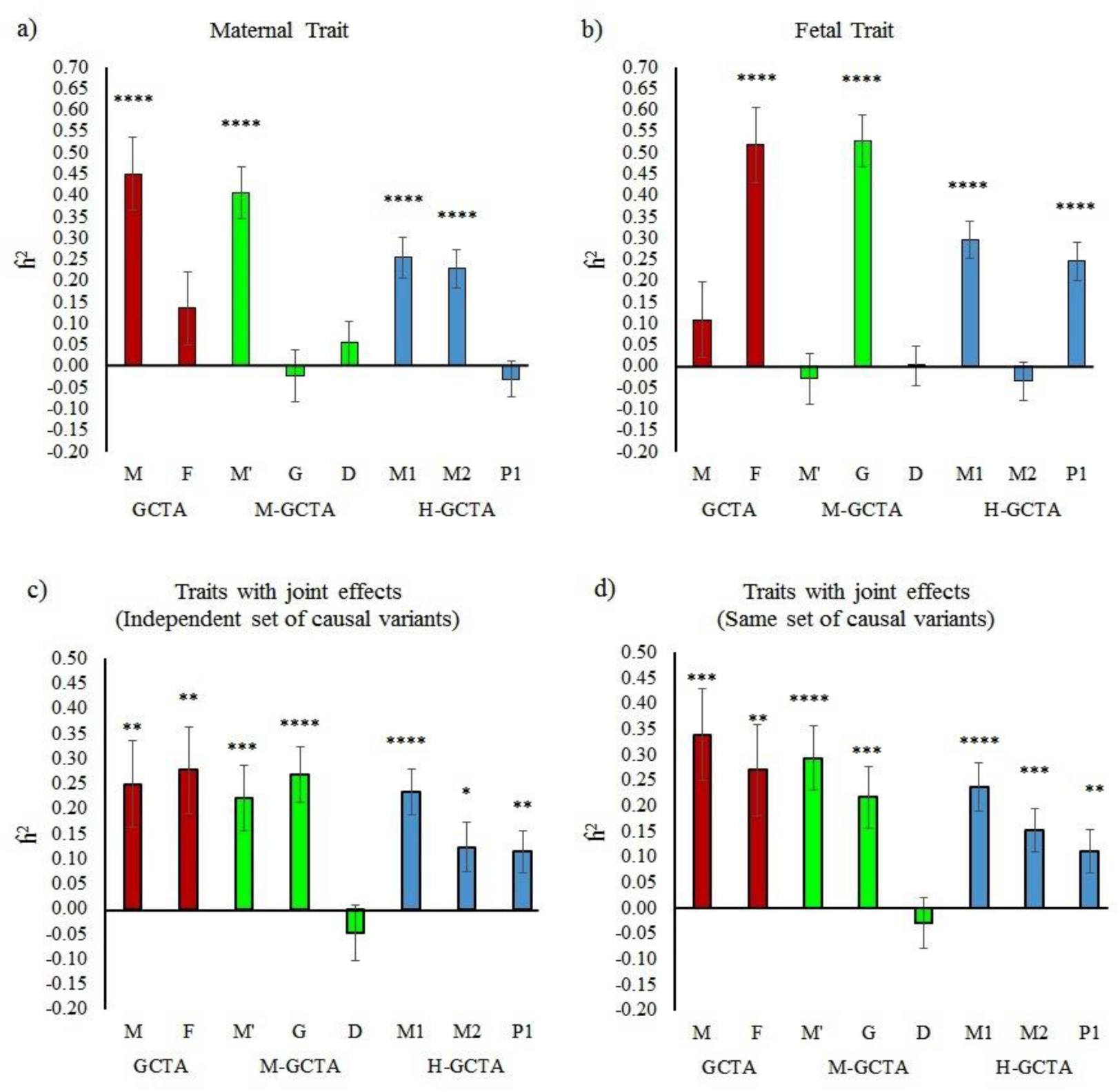
Comparison of 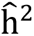 for simulated traits from pooled dataset, estimated through different approaches fitting GREML (α = –1.0): a) maternal traits; b) fetal traits; c) traits where independent sets of causal variants have effects through mother and fetus; d) traits where same set of causal variants have effects through mother and fetus. For GCTA, M is the GRM generated from maternal genotypes (m), and F is the GRM generated from fetal genotypes (f). For M-GCTA, M’ represents the genetic relationship matrix of mothers; G represents genetic relationship matrix of children and D represents mother-child covariance matrix. For H-GCTA, M1 is the GRM generated from maternal transmitted alleles (m1), M2 is the GRM generated from maternal non-transmitted alleles (m2), and P1 is the GRM generated from paternal transmitted alleles (p1). A total of 100 replicates of each phenotype were simulated using empirical genotypes of <u>Pooled</u> dataset. P-values were calculated using z test statistics (two sided). * = (p value < 5.0E-02), ** = (p value < 1.0E-02), *** = (p value < 1.0E-03) and **** = (p value < 1.0E-04).

### Heritability of simulated traits with only fetal effects

Like maternal traits, we used conventional GCTA to estimate 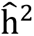 for fetal traits in mothers and children separately. The estimated 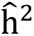 based on m and f were 10.9% (S.E. = 8.8%) and 51.9% (S.E. = 8.8%) respectively (Figure 2b, Supplementary Table 14). Similarly, using M-GCTA for fetal traits in mother-child duos, variance attributable to indirect maternal effect (M’), direct fetal effect (G) and direct-indirect effect covariance (D) were –3.0% (S.E. = 6.0%), 52.7% (S.E. = 6.1%) and 0.0% (S.E. = 4.7%) respectively (Figure 2b, Supplementary Table 14). Using H-GCTA in mother-child duos, we estimated the variance of the simulated fetal traits attributable to m1 (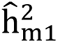 = 29.6%, S. E. = 4.3%), m2 (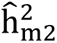 = −3.6%, S. E. = 4.4%) and p1 (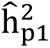 = 24.6%, S. E. = 4.4%) (Figure 2b, Supplementary Table 14). While, conventional GCTA estimated superficial contributions from maternal genotypes besides fetal genotypes, M-GCTA and H-GCTA clearly showed the fetal origin of the simulated phenotypes. As compared to M-GCTA, H-GCTA further resolved almost equal contributions from maternal and paternal transmitted alleles through m1 and p1.

### Heritability of simulated traits with independent maternal-fetal genetic effects

Traits with independent maternal and fetal effects were simulated in two ways – using same set and different sets of causal variants in mothers and children. Using independent sets of causal variants and conventional GCTA approach, the estimated 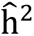 based on m and f were 24.9% (S.E. = 8.7%) and 27.8% (S.E. = 8.7%) respectively (Figure 2c; Supplementary Table 15). Using M-GCTA approach, variance attributable to indirect maternal effect (M’), direct fetal effect (G) and direct-indirect effect covariance (D) were estimated as 22.1% (S.E. = 6.6%), 26.8% (S.E. = 5.6%) and –4.7% (S.E. = 5.5%) respectively (Figure 2c, Supplementary Table 15). Conversely, H-GCTA estimated the genetic variance attributable to m1 (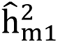 = 23.4%, S. E. = 4.6%), m2 (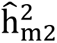 = 12.4%, S. E. = 4.8%) and p1 (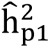 = 11.5%, S. E. = 4.2%) (Figure 2c, Supplementary Table 15). We observed similar results from traits, simulated using same set of causal variants with independent maternal-fetal genetic effects in mothers and children (Figure 2d, Supplementary Table 16). We observed that conventional GCTA and M-GCTA showed equal contribution of maternal and fetal genotypes to the phenotypic variance of the simulated phenotypes. As compared to M-GCTA, H-GCTA estimated the contributions of maternal transmitted (m1), maternal non-transmitted (m2) and paternal transmitted alleles (p1) as expected i.e. 2:1:1 (Supplementary Figure 5, Supplementary Tables 15 and 16) which demonstrated equal and independent maternal and fetal contributions.

### Heritability of simulated traits with correlated maternal-fetal genetic effects

We simulated traits influenced by joint maternal-fetal genetic effects with average negative (–0.5, –1.0) and positive (0.5, 1.0) correlation by using same set of causal variants in mothers and children. For traits with 100% negative correlation of maternal-fetal genetic effects, the estimated 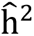 using conventional GCTA approach, were 8.2% (S.E. = 9.4%) and 11.4% (S.E. = 9.4%) based on m and f, respectively (Figure 3a and Supplementary Table 17). Using M-GCTA approach, the variance attributable to indirect maternal effect (M’), direct fetal effect (G) and direct-indirect effect covariance (D) were estimated as 35.9% (S.E. = 8.4%), 38.3% (S.E. = 6.8%) and –36.1% (S.E. = 6.2%) respectively (Figure 3a and Supplementary Table 17). H-GCTA further partitioned the variance into variance components attributable to m1 (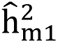 = −1.0%, S. E. = 5.0%), m2 (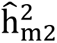 = 17.6%, S. E. = 5.2%) and p1 (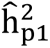 = 20.1%, S. E. = 4.1%) (Figure 3a and Supplementary Table 17). Although, negative values of 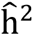 usually mean nothing and are considered as zero, they are important for interpretation of results for traits with negative correlation of maternal-fetal genetic effects. We observed that conventional GCTA substantially underestimated the genetic contribution of maternal and fetal genomes. As expected, M-GCTA estimated equal contribution of maternal and fetal genetic effects to the phenotypic variance whereas negative and equal contribution of direct-indirect effect covariance suggested 100% negative correlation of maternal-fetal genetic effects. Similarly, H-GCTA showed no contribution from m1 (due to 100% negative correlation of maternal-fetal genetic effects) and almost equal contribution from m2 and p1 to the phenotypic variance (Supplementary Figure 5). We observed similar patterns for traits with 50% negative correlation of maternal-fetal genetic effects (Figure 3b, Supplementary Table 18).

**Figure 3:**
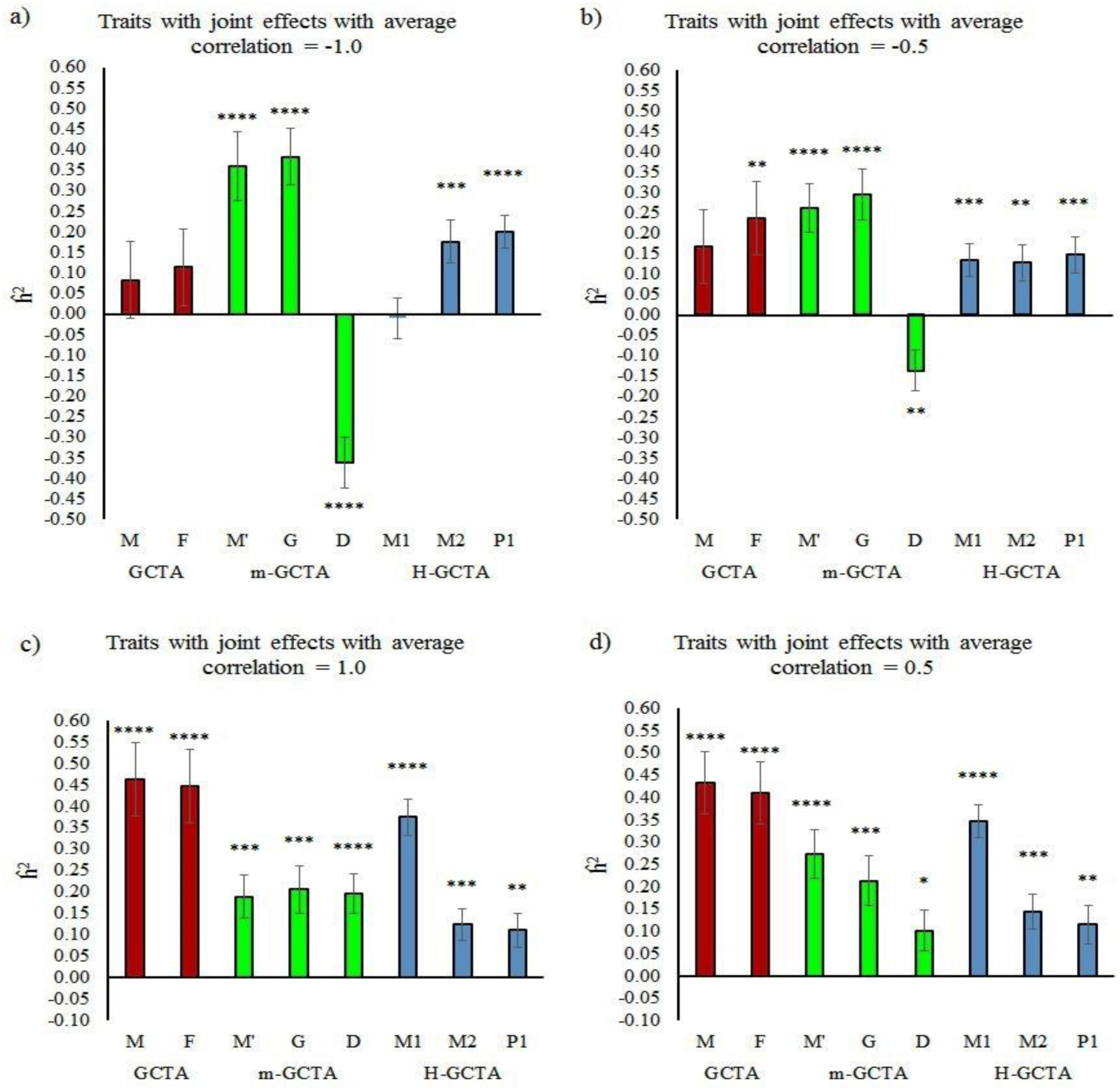
Comparison of 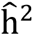 estimated through different approaches fitting GREML (α = –1.0) for simulated traits with joint maternal–fetal effects from pooled dataset: a) average correlation = –1.0; b) average correlation = –0.5; c) average correlation = 1.0; d) average correlation = 0.5. For GCTA, M is the GRM generated from maternal genotypes (m), and F is the GRM generated from fetal genotypes (f). For M-GCTA, M’ represents the genetic relationship matrix of mothers; G represents genetic relationship matrix of children and D represents mother-child covariance matrix. For H-GCTA, M1 is the GRM generated from maternal transmitted alleles (m1), M2 is the GRM generated from maternal non-transmitted alleles (m2), and P1 is the GRM generated from paternal transmitted alleles (p1). A total of 100 replicates of each phenotype were simulated using empirical genotypes of Pooled dataset. P-values were calculated using z test statistics (two sided). * = (p value < 5.0E-02), ** = (p value < 1.0E-02), *** = (p value < 1.0E-03) and **** = (p value < 1.0E-04)

Similarly, conventional GCTA approach estimated 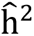 based on m and f as 46.2% (S.E. = 8.6%) and 44.8% (S.E. = 8.6%), respectively for traits with 100% positive correlation of maternal-fetal genetic effects (Figure 3c, Supplementary Table 19). M-GCTA approach estimated the variance attributable to indirect maternal effect (M’), direct fetal effect (G) and direct-indirect effect covariance (D) as 18.9% (S.E. = 5.0%), 20.6% (S.E. = 5.6%) and 19.7% (S.E. = 4.6%), respectively (Figure 3c, Supplementary Table 19). Using H-GCTA, we estimated the genetic variance of simulated traits attributable to m1, m2 and p1 [(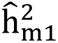 = 37.4%, S. E. = 4.2%), (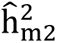 = 11.4%, S. E. = 3.7%), (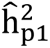 = 11.1%, S. E. = 3.9%)] (Figure 3c, Supplementary Table 19). While conventional GCTA substantially overestimated the variance attributable to maternal and fetal genotypes, M-GCTA estimated equal contribution of indirect maternal effects, direct fetal effects and direct-indirect effects covariance to the phenotypic variance. Similarly, H-GCTA showed much larger contribution from m1 and equal contribution from m2 and p1 to the phenotypic variance which follows a ratio of 4:1:1 in case of 100% positive correlation of maternal-fetal genetic effects (Supplementary Figure 5). Similar patterns were observed for traits with 50% positive correlation of maternal-fetal genetic effects (Figure 3d, Supplementary Table 20).

### Heritability of simulated fetal traits with POEs

We also estimated genetic variance using GREML for simulated fetal traits with different levels of parent-of-origin effects (POEs) in varying proportion of causal variants. We simulated two scenarios where maternal imprinting was mimicked by reducing the effect of m1 as compared to p1 in 25% and 50% of the causal variants. In each scenario, we generated a range of imprinting patterns such as u_m1_⁄u_p1_ = 0.75, u_m1_⁄u_p1_ = 0.50, u_m1_⁄u_p1_ = 0.25 and u_m1_⁄u_p1_ = 0. The first three conditions represented partial maternal imprinting whereas the last condition i.e. u_m1_⁄u_p1_ = 0 represented complete maternal imprinting. Using our approach (H-GCTA), we estimated the total 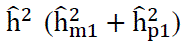 as expected (∼ 50%) (Figure 4, Supplementary Table 21). Results from H-GCTA showed that the variance attributable to m1 (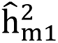) decreased whereas the variance attributable to p1 (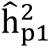) increased in accordance with the level of imprinting in each scenario (Figure 4a and 4b). We also compared results from our approach with those from GCTA and M-GCTA. While GCTA and M-GCTA were unable to detect contribution of parent-of-origin effects (POEs), H-GCTA detected the variance attributable to POEs as 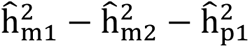 (Supplementary Table 21).

**Figure 4:**
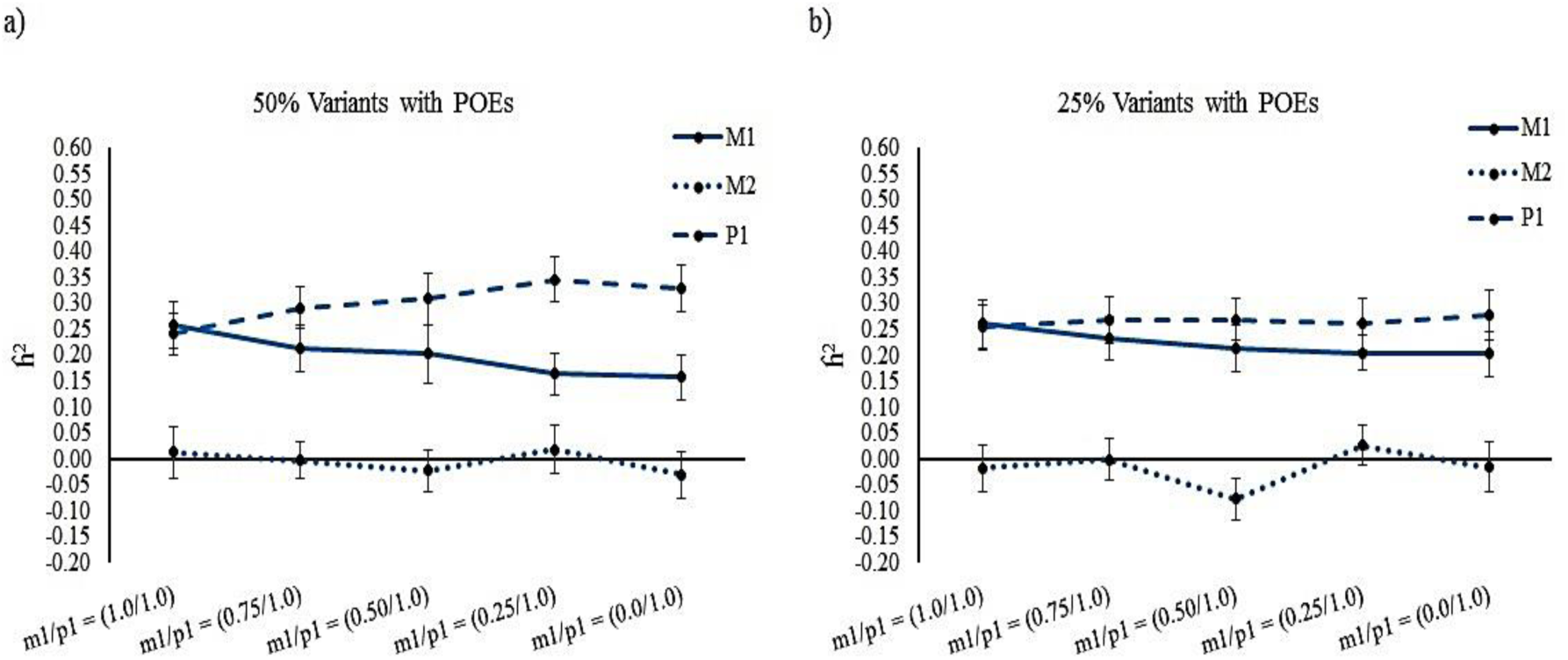
Variance attributable to m1, m2 and p1 estimated through H-GCTA using GREML (α = –1.0) model in simulated fetal traits from pooled dataset: a) 50% causal variants with POEs; b) 25% causal variants with POEs. POEs were incorporated by reducing the effect of m1 as compared to p1 by multiplying effects of m1 with (1 – I) where I is the imprinting factor such as 0.25, 0.50, 0.75 and 1.0. In each scenario, m1 shows either no imprinting i.e. I = 0.0 (m1⁄p1 = 1.0⁄1.0) or partial imprinting i.e. I = 0.25-0.75 (m1⁄p1 = 0.75⁄1.0 − 0.25⁄1.0) or complete imprinting i.e. I = 1.0 (m1⁄p1 = 0.0⁄1.0).

### Heritability estimation of pregnancy-related outcomes using empirical data

All analyses for the estimation of genetic variance were performed using imputed genotype data of ∼ 11 million markers across 10,375 mother-child pairs. In addition, two MAF cut-offs (0.001 and 0.01) yielding approximately 9 million and 7 million markers respectively, were used for analysis. Only independent mother-child pairs (kinship coefficient < 0.05) were used in analysis and 20 principal components (PCs) were used along with genotype-based GRMs in LMM (Supplementary Figure 6). For haplotype-based GRMs, we used 30 PCs (10 PCs corresponding to each haplotype) as covariates in LMM (Supplementary Figure 6). Like simulated traits, we estimated genetic variance using three approaches – conventional GCTA approach, M-GCTA approach and H-GCTA approach. For each approach, we fitted three models – GREML, LDAK-Thin and LDAK-Weights. Two values of α (–0.25 and –1.0), which represents the extent to which minor allele frequency (MAF) influences the variance of SNP effects on phenotypes were used for each model. Here, we describe results based on GRMs calculated through all polymorphic SNPs and three models with recommended α values i.e. GREML (α = –1.0), LDAK-Thin (α = –0.25) and LDAK-Weights (α = –0.25). Results based on all polymorphic SNPs using other models are provided in supplementary table 22. Similarly, results based on GRMs calculated through SNPs with MAF > 0.001 and SNPs with MAF > 0.01 are provided in supplementary text, supplementary table 23 and 24.

### Heritability of gestational duration

Using GREML (α = –1.0), the conventional GCTA approach estimated 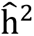 of gestational duration based on m and f – (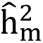 = 31.4%; S.E. = 5.4%) and (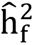 = 12.2%; S.E. = 5.2%). Our approach (H-GCTA) further resolved the variance attributable to m1 – 17.3% (S.E. = 5.2%;), m2 – 12.3% (S.E. = 5.2%) and p1 – 0.0% (S.E. = 5.0%) (Figure 5a, Table 2a). Results using our approach suggested that the genetic variance in gestational duration was primarily influenced by maternal genome i.e. the SNPs which influence gestational duration through maternal genetic effect. Comparison with M-GCTA confirmed the results from H-GCTA (Figure 5a, Table 2a). The genetic variance estimated through LDAK-Thin (α = –0.25) was similar to those obtained from GREML (α = –1.0). However, estimates from LDAK-Weights (α = –0.25) were substantially larger than those obtained from GREML (α = –1.0) (Table 2a). This pattern is consistent with previous observation for other traits using LDAK model^16^ and thoroughly discussed elsewhere^13^

**Figure 5:**
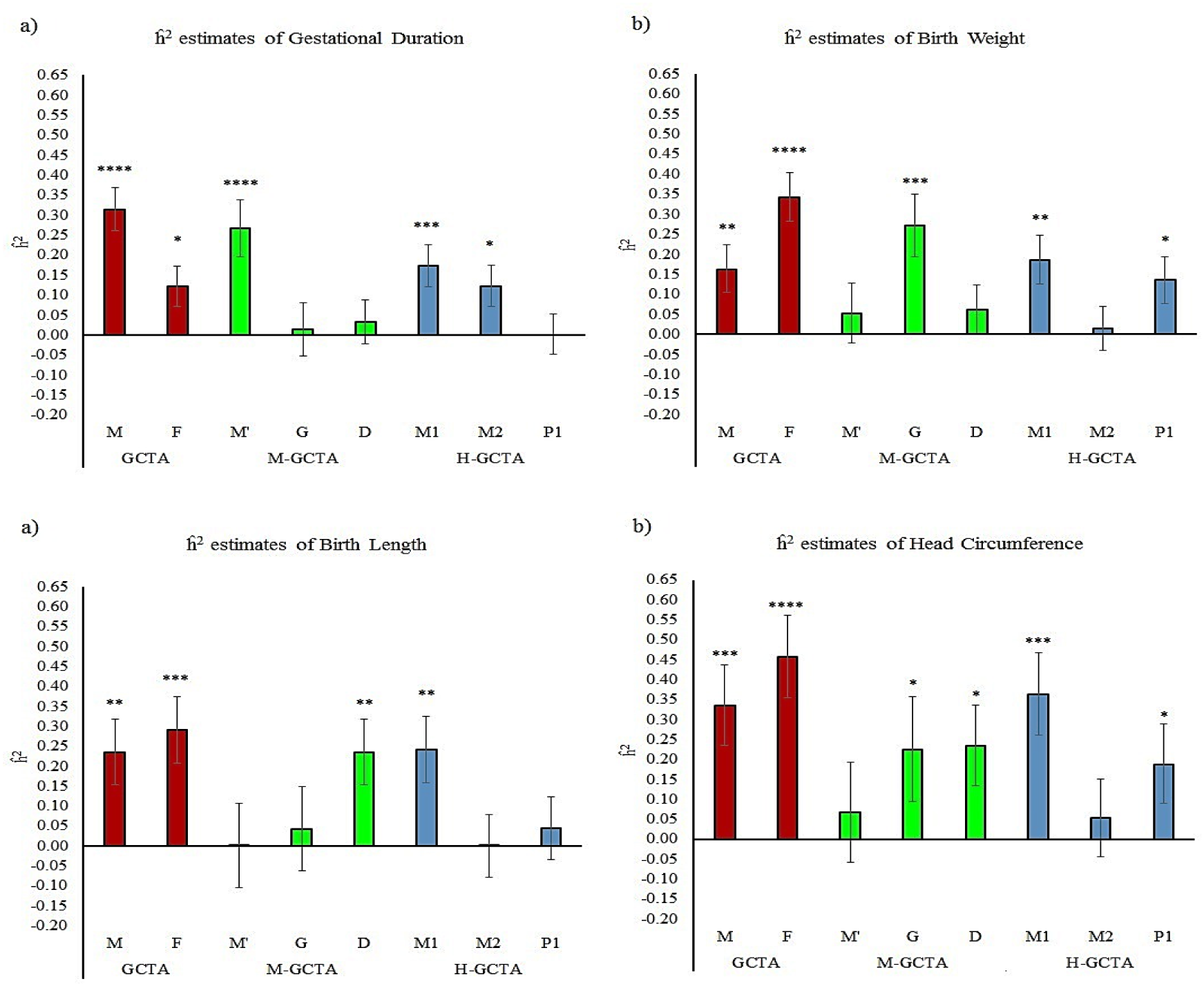
Comparison of 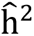 estimated through different approaches fitting GREML (α = –1.0) for pregnancy-related outcomes in unrelated mother-child pairs (relatedness cutoff > 0.05): a) gestational duration, b) birth weight, c) birth length, and d) head circumference. For GCTA, M is the GRM generated from maternal genotypes (m), and F is the GRM generated from fetal genotypes (f). For M-GCTA, M’ represents the genetic relationship matrix of mothers; G represents genetic relationship matrix of children and D represents mother-child covariance matrix. For H-GCTA, M1 is the GRM generated from maternal transmitted alleles (m1), M2 is the GRM generated from maternal non-transmitted alleles (m2), and P1 is the GRM generated from paternal transmitted alleles (p1). For conventional GCTA and M-GCTA approach analyses were adjusted for 20 principal components (PCs) whereas for H-GCTA, analyses were adjusted for 30 PCs (10 PCs corresponding to m1, m2 and p1 each). P-values were calculated using z test statistics (one sided). * = (p value < 5.0E-02), ** = (p value < 1.0E-02), *** = (p value < 1.0E-03) and **** = (p value < 1.0E-04)

**Table 2:**
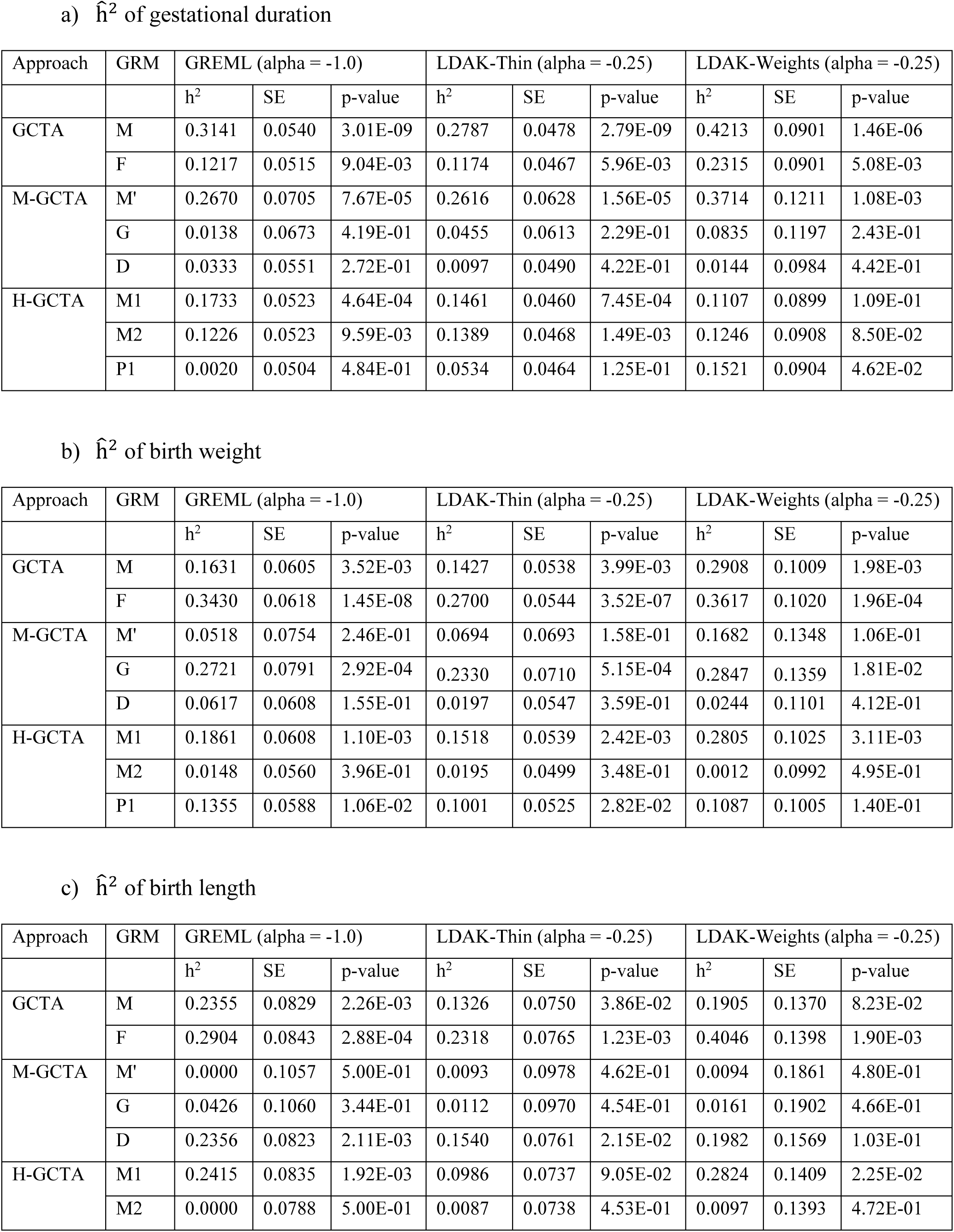
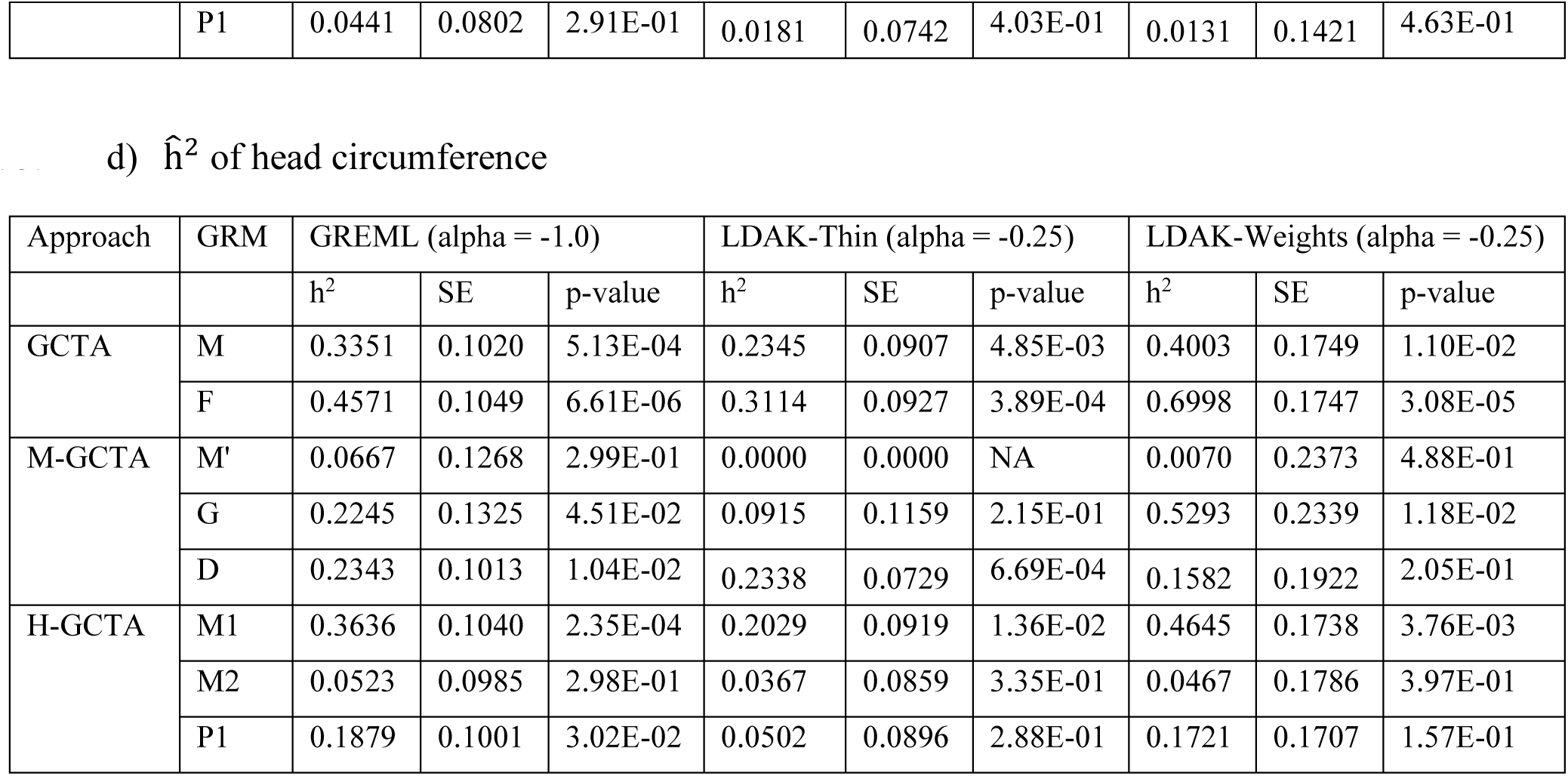
Comparison of 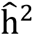 estimated through different approaches fitting GREML (α = –1.0), LDAK-Thin (α = –0.25) and LDAK-Weights (α = –0.25) for a) gestational duration, b) birth weight, c) birth length and d) head circumference. GRMs were generated using all polymorphic SNPs. For GCTA, M is the GRM generated from maternal genotypes (m), and F is the GRM generated from fetal genotypes (f). For M-GCTA, M’ represents the genetic relationship matrix of mothers; G represents genetic relationship matrix of children and D represents mother-child covariance matrix. For H-GCTA, M1 is the GRM generated from maternal transmitted alleles (m1), M2 is the GRM generated from maternal non-transmitted alleles (m2), and P1 is the GRM generated from paternal transmitted alleles (p1). Gestational duration was adjusted for fetal sex and fetal size measurements at birth were additionally adjusted for gestational duration up to third orthogonal polynomial. Analyses using GCTA and M-GCTA approach were adjusted for 20 PCs and H-GCTA approach was adjusted for 30 PCs (10 PCs corresponding to m1, m2 and p1 each). P-values were calculated using z test statistics (one sided).

| Approach | GRM | GREML (alpha = -1.0) |  |  | LDAK-Thin (alpha = -0.25) |  |  | LDAK-Weights (alpha = -0.25) |  |  |
| --- | --- | --- | --- | --- | --- | --- | --- | --- | --- | --- |
| | | $h^2$ | SE | p-value | $h^2$ | SE | p-value | $h^2$ | SE | p-value |
| GCTA | M | 0.3141 | 0.0540 | 3.01E-09 | 0.2787 | 0.0478 | 2.79E-09 | 0.4213 | 0.0901 | 1.46E-06 |
|  | F | 0.1217 | 0.0515 | 9.04E-03 | 0.1174 | 0.0467 | 5.96E-03 | 0.2315 | 0.0901 | 5.08E-03 |
| M-GCTA | M' | 0.2670 | 0.0705 | 7.67E-05 | 0.2616 | 0.0628 | 1.56E-05 | 0.3714 | 0.1211 | 1.08E-03 |
|  | G | 0.0138 | 0.0673 | 4.19E-01 | 0.0455 | 0.0613 | 2.29E-01 | 0.0835 | 0.1197 | 2.43E-01 |
|  | D | 0.0333 | 0.0551 | 2.72E-01 | 0.0097 | 0.0490 | 4.22E-01 | 0.0144 | 0.0984 | 4.42E-01 |
| H-GCTA | M1 | 0.1733 | 0.0523 | 4.64E-04 | 0.1461 | 0.0460 | 7.45E-04 | 0.1107 | 0.0899 | 1.09E-01 |
|  | M2 | 0.1226 | 0.0523 | 9.59E-03 | 0.1389 | 0.0468 | 1.49E-03 | 0.1246 | 0.0908 | 8.50E-02 |
|  | P1 | 0.0020 | 0.0504 | 4.84E-01 | 0.0534 | 0.0464 | 1.25E-01 | 0.1521 | 0.0904 | 4.62E-02 |

b) $\hat{h}^2$ of birth weight
| Approach | GRM | GREML (alpha = -1.0) |  |  | LDAK-Thin (alpha = -0.25) |  |  | LDAK-Weights (alpha = -0.25) |  |  |
| --- | --- | --- | --- | --- | --- | --- | --- | --- | --- | --- |
| | | $h^2$ | SE | p-value | $h^2$ | SE | p-value | $h^2$ | SE | p-value |
| GCTA | M | 0.1631 | 0.0605 | 3.52E-03 | 0.1427 | 0.0538 | 3.99E-03 | 0.2908 | 0.1009 | 1.98E-03 |
|  | F | 0.3430 | 0.0618 | 1.45E-08 | 0.2700 | 0.0544 | 3.52E-07 | 0.3617 | 0.1020 | 1.96E-04 |
| M-GCTA | M' | 0.0518 | 0.0754 | 2.46E-01 | 0.0694 | 0.0693 | 1.58E-01 | 0.1682 | 0.1348 | 1.06E-01 |
|  | G | 0.2721 | 0.0791 | 2.92E-04 | 0.2330 | 0.0710 | 5.15E-04 | 0.2847 | 0.1359 | 1.81E-02 |
|  | D | 0.0617 | 0.0608 | 1.55E-01 | 0.0197 | 0.0547 | 3.59E-01 | 0.0244 | 0.1101 | 4.12E-01 |
| H-GCTA | M1 | 0.1861 | 0.0608 | 1.10E-03 | 0.1518 | 0.0539 | 2.42E-03 | 0.2805 | 0.1025 | 3.11E-03 |
|  | M2 | 0.0148 | 0.0560 | 3.96E-01 | 0.0195 | 0.0499 | 3.48E-01 | 0.0012 | 0.0992 | 4.95E-01 |
|  | P1 | 0.1355 | 0.0588 | 1.06E-02 | 0.1001 | 0.0525 | 2.82E-02 | 0.1087 | 0.1005 | 1.40E-01 |

c) $\hat{h}^2$ of birth length
| Approach | GRM | GREML (alpha = -1.0) |  |  | LDAK-Thin (alpha = -0.25) |  |  | LDAK-Weights (alpha = -0.25) |  |  |
| --- | --- | --- | --- | --- | --- | --- | --- | --- | --- | --- |
| | | $h^2$ | SE | p-value | $h^2$ | SE | p-value | $h^2$ | SE | p-value |
| GCTA | M | 0.2355 | 0.0829 | 2.26E-03 | 0.1326 | 0.0750 | 3.86E-02 | 0.1905 | 0.1370 | 8.23E-02 |
|  | F | 0.2904 | 0.0843 | 2.88E-04 | 0.2318 | 0.0765 | 1.23E-03 | 0.4046 | 0.1398 | 1.90E-03 |
| M-GCTA | M' | 0.0000 | 0.1057 | 5.00E-01 | 0.0093 | 0.0978 | 4.62E-01 | 0.0094 | 0.1861 | 4.80E-01 |
|  | G | 0.0426 | 0.1060 | 3.44E-01 | 0.0112 | 0.0970 | 4.54E-01 | 0.0161 | 0.1902 | 4.66E-01 |
|  | D | 0.2356 | 0.0823 | 2.11E-03 | 0.1540 | 0.0761 | 2.15E-02 | 0.1982 | 0.1569 | 1.03E-01 |
| H-GCTA | M1 | 0.2415 | 0.0835 | 1.92E-03 | 0.0986 | 0.0737 | 9.05E-02 | 0.2824 | 0.1409 | 2.25E-02 |
|  | M2 | 0.0000 | 0.0788 | 5.00E-01 | 0.0087 | 0.0738 | 4.53E-01 | 0.0097 | 0.1393 | 4.72E-01 |
|  | P1 | 0.0441 | 0.0802 | 2.91E-01 | 0.0181 | 0.0742 | 4.03E-01 | 0.0131 | 0.1421 | 4.63E-01 |

**Table 2:** Comparison of $\hat{h}^2$ estimated through different approaches fitting GREML ( $\alpha = -1.0$ ), LDAK-Thin ( $\alpha = -0.25$ ) and LDAK-Weights ( $\alpha = -0.25$ ) for a) gestational duration, b) birth weight, c) birth length and d) head circumference.
| Approach | GRM | GREML ($\alpha = -1.0$) | | | LDAK-Thin ($\alpha = -0.25$) | | | LDAK-Weights ($\alpha = -0.25$) | | |
| --- | --- | --- | --- | --- | --- | --- | --- | --- | --- | --- |
| | | $h^2$ | SE | p-value | $h^2$ | SE | p-value | $h^2$ | SE | p-value |
| GCTA | M | 0.3351 | 0.1020 | 5.13E-04 | 0.2345 | 0.0907 | 4.85E-03 | 0.4003 | 0.1749 | 1.10E-02 |
|  | F | 0.4571 | 0.1049 | 6.61E-06 | 0.3114 | 0.0927 | 3.89E-04 | 0.6998 | 0.1747 | 3.08E-05 |
| M-GCTA | M' | 0.0667 | 0.1268 | 2.99E-01 | 0.0000 | 0.0000 | NA | 0.0070 | 0.2373 | 4.88E-01 |
|  | G | 0.2245 | 0.1325 | 4.51E-02 | 0.0915 | 0.1159 | 2.15E-01 | 0.5293 | 0.2339 | 1.18E-02 |
|  | D | 0.2343 | 0.1013 | 1.04E-02 | 0.2338 | 0.0729 | 6.69E-04 | 0.1582 | 0.1922 | 2.05E-01 |
| H-GCTA | M1 | 0.3636 | 0.1040 | 2.35E-04 | 0.2029 | 0.0919 | 1.36E-02 | 0.4645 | 0.1738 | 3.76E-03 |
|  | M2 | 0.0523 | 0.0985 | 2.98E-01 | 0.0367 | 0.0859 | 3.35E-01 | 0.0467 | 0.1786 | 3.97E-01 |
|  | P1 | 0.1879 | 0.1001 | 3.02E-02 | 0.0502 | 0.0896 | 2.88E-01 | 0.1721 | 0.1707 | 1.57E-01 |

### Heritability of gestational duration adjusted birth weight

Analysis using conventional GCTA showed that the estimated 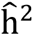 of birth weight based on m and f were 16.3% (S.E. = 6.1%) and 34.3% (S.E. = 6.2%) respectively. Using our approach, we further distinguished the variance attributable to m1 – 18.6% (S.E. = 6.1%); m2 – 1.5% (S.E. = 5.6%) and p1 – 13.6% (S.E. = 5.9%) (Figure 5b, Table 2b). The estimates obtained through H-GCTA suggested that genetic variance in birth weight was primarily determined by the fetal genome. Comparison of genetic variance estimated from our approach with those from M-GCTA illustrated that genetic variance in birth weight was mainly attributable to the SNPs which influence birth weight only through direct fetal effect (Figure 5b, Table 2b). Like gestational duration, genetic variance estimated through LDAK-Thin (α = –0.25) was similar to those obtained from GREML (α = –1.0) whereas LDAK-Weights (α = –0.25) estimated larger 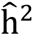.Heritability of gestational duration adjusted birth length

### Heritability of gestational duration adjusted birth length

We estimated 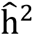 of birth length based on m (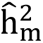 = 23.6%; S.E. = 8.3%) and f (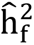 = 29.0%; S.E. = 8.4%) using conventional GCTA approach. While M-GCTA indicated that birth length is largely influenced by positive maternal-fetal covariance (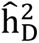 = 23.6%; S.E. = 8.2%), H-GCTA resolved the genetic variance attributable to m1 – 24.2% (S.E. = 8.4%); m2 – 0.0% (S.E. = 7.9%) and p1 – 4.4% (S.E. = 8.0%) (Figure 5c, Table 2c). H-GCTA showed that unlike birth weight, variance in birth length was mainly attributable to m1 with a much smaller attribution to p1 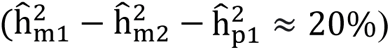. According to our simulations, this pattern i.e. 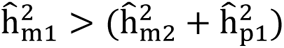 could be generated due to either positively correlated maternal-fetal genetic effects (Figure 3, Supplementary Tables 19, 20) or POEs (Figure 4, Supplementary Table 21). A previous study using M-GCTA^21^ suggested that birth length was influenced by both maternal and fetal genome with different genes contributing to the maternal and fetal effects. However, current study indicates that variance in birth length is primarily attributable to positively correlated maternal-fetal genetic effects along with possible POEs 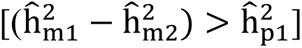.

### Heritability of gestational duration adjusted head circumference

SNP-based narrow-sense heritability (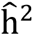) of head circumference estimated using a conventional GCTA approach was 33.5% (S.E. = 10.2%) and 45.7% (S.E. = 10.5%) based on m and f, respectively. Using H-GCTA, we resolved the variance attributable to maternal and fetal genomes into m1 – 36.4% (S.E. = 10.4%); m2 – 5.2% (S.E. = 9.9%) and p1 – 18.8% (S.E. = 10.0%) (Figure 5d, Table 2d). The difference between 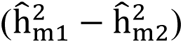 and 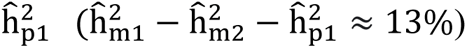 suggested that head circumference is largely influenced by fetal genetic effects along with either correlated maternal-fetal genetic effects or possible POEs or both. Similarly, the results from M-GCTA analysis showed approximately equal contribution to variance of head circumference from G and D (Table 2d). The comparison of results from H-GCTA and M-GCTA suggested that head circumference was primarily determined by fetal genome i.e. phenotypic variance of head circumference was largely influenced by direct fetal effects along with positively correlated joint maternal-fetal effects or POEs. The results also suggested some influence through explicit maternal genetic effect (Table 2d).

## Discussion

Unlike widely studied complex human traits^8,9,13,16,33^, pregnancy-related outcomes are simultaneously influenced by maternal and fetal genomes. Therefore, conventional genotype-based approaches that were developed to estimate the genetic contribution to phenotypic variance are limited in addressing the confounding of shared alleles between maternal and fetal genomes. Here, we consider the mother-child pair as a single analytical unit with three haplotypes – maternal transmitted (m1), maternal non-transmitted (m2) and paternal transmitted (p1). Using such an analytical unit, we simultaneously disentangle the contribution of m1 (exclusive and joint maternal-fetal effects), m2 (exclusive maternal effect) and p1 (exclusive fetal effect) to the phenotypic variance. Using the simulated data with varying contributions and correlation of maternal and fetal genetic effects, we show that our newly developed H-GCTA approach can explicitly resolve maternal and fetal contributions and outperforms the GCTA and M-GCTA approach, particularly in the presence of POEs (Figure 4, Supplementary Table 21). We further apply our haplotype-based approach to distinguish the genetic contribution of mothers and offspring to the phenotypic variance of gestational duration and gestational duration adjusted fetal size measurements at birth in 10,375 European mother-child pairs. A comparison of results from H-GCTA with those from M-GCTA and conventional GCTA approach reveals that gestational duration is primarily influenced by maternal genome whereas fetal size measurements at birth are largely driven by fetal genome. The new results not only confirm the previous findings from epidemiological^34–40^ and genetic^21–25,27,31,41–46^ studies but also provide new insights into the genetic architecture of fetal size at birth.

The results based on ∼11 million polymorphic SNPs show that approximately 17% and 12% variance in gestational duration is attributable to the m1 and m2, respectively with a minimal contribution from p1 (Figure 5; Table 2). In contrast, variance in gestational duration adjusted fetal size measurements at birth are mainly contributed by m1 (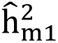 = 19-36%) and p1 (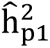 = 4-14%) with a minimal contribution from m2 (Figure 5; Table 2). Among fetal size measurements at birth, variance in birth weight has significant contributions from m1 (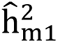 = 19%) as well as p1 (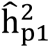 = 14%) whereas variance in birth length and head circumference are mainly attributable to m1 (birth length: 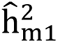 = 24%; head circumference: 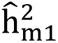 = 36%). These new results suggest that variance in gestational duration is mainly attributable to indirect maternal genetic effects whereas variance in birth weight is mainly attributable to direct fetal genetic effects. In addition, a larger contribution of m1 as compared to m2 and p1 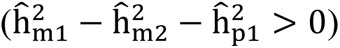 to the variance of birth length and head circumference suggests a substantial contribution of correlated maternal-fetal genetic effects or possible POEs or both (Table 2). Results using SNPs with MAF > 0.001 and SNPs with MAF > 0.01 showed similar results (Supplementary Tables 23 and 24).

As observed in the analyses of simulated traits, estimated genetic variance observed through GREML (α = –1.0) and LDAK-Thin (α = –0.25) are similar for pregnancy-related outcomes. Consistent with previous reports^13,16^, estimated genetic variance using LDAK-Weights (α = –0.25) are up to 30% higher than those using GREML (α = –1.0). However, analysis using LDAK-Thin (α = –0.25) provides slightly lower estimates for gestational duration and birth weight and substantially lower estimates for birth length and head circumference. Similarly, GREML (α = – 0.25) and LDAK-Weights (α = –1.0) estimate substantially smaller 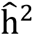 whereas LDAK-Thin (α = – 1.0) estimates substantially larger 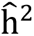 (Supplementary Table 22). These results are consistent with the results of simulated traits in the current study and could be due to misspecification of analytical model^47^. For GREML (α = –1.0) model, we observe the largest estimates of genetic variance for each trait using all polymorphic SNPs, which decreases with increasing threshold of MAF cutoff (number of SNPs decrease with increasing MAF cutoff) (Supplementary Tables 23 and 24). The decrease in the estimated genetic variance with decrease in number of markers is a general limitation of GREML model which is dependent on several assumptions^13,48^.

In general, results for pregnancy-related outcomes follow a similar pattern as those for simulated traits. Specifically, estimated genetic variance of gestational duration and birth weight mimic a pattern similar to the simulated maternal and fetal traits, respectively. Interestingly, estimated genetic variance of head circumference follow a mixed pattern with a large fetal and small maternal genetic influence along with a large influence of maternal transmitted alleles. Irrespective of the analytical models and MAF cut-offs for GRM calculation, H-GCTA estimates a larger contribution of m1 as compared to p1 with almost no contribution of m2 to the phenotypic variance of birth length (Table 2, Supplementary Tables 23 and 24). Similarly, M-GCTA estimates a larger contribution of correlated maternal-fetal genetic effects (D) as compared to direct fetal effects (G) with almost no contribution of indirect maternal effect (M’) (Table 2, Supplementary Tables 23 and 24). It is possible that there could be complicated maternal-fetal interactions that are not modeled by any of these approaches.

Interestingly, we observe that the contribution of m1 is larger than m2 or p1 for every pregnancy phenotype in the current study. There are several possible explanations for this pattern of results. The most obvious explanation is that m1 can influence a pregnancy phenotype through both the mother and fetus. For example, for a trait mainly defined by the maternal genome like gestational duration, higher contribution of m1 in comparison to m2 could be due to small but non-zero fetal effect of the m1 alleles. Similarly, for traits mainly defined by the fetal genome such as fetal size measurements at birth, higher contribution of m1 in comparison to p1 could be due to small but non-zero maternal effect of the m1 alleles. Assuming maternal-fetal additivity (genetic effects through mother and fetus influence a pregnancy-related outcome in additive manner) with independent maternal-fetal genetic effects and no POEs, 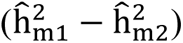 is equal to 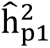. A larger value of 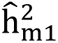 as compared to 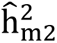 and 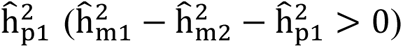 suggests presence of either positively correlated maternal-fetal genetic effects or possible POEs or both. For birth length and head circumference, the near zero maternal genetic effect (as estimated by M’ in the M-GCTA analysis) and the null contribution from the maternal non-transmitted alleles (m2) suggests possible existence of POEs along with positively correlated maternal-fetal genetic effects. Besides the above-mentioned explanations, several other biological phenomena such as interaction between SNPs within the mother or fetus (epistasis) and gene-environment interaction may influence the pattern of genetic variance of pregnancy outcomes.

Despite the above advances, our current approach has certain limitations. Our approach by itself cannot explicitly distinguish the contribution of correlated maternal-fetal genetic effects from POEs. Current haplotype-based approach attempts to relax some of the underlying assumptions in conventional and contemporary approaches such as equal effects of maternal and paternal transmitted alleles in fetus and allelic additivity. However, the interpretation of the results requires assumptions on maternal-fetal additivity and random mating population. In addition, heritability estimation in our approach can also be affected if assumptions such as absence of epistasis (gene-gene interaction) and gene-environment interaction are not met.

In conclusion, we introduce an approach (H-GCTA) to partition phenotypic variance of pregnancy outcomes to maternal transmitted, non-transmitted and paternal transmitted alleles in mother/child pairs. This method provides a direct way to dissect the maternal and fetal genetic contributions to pregnancy-related outcomes. In addition, H-GCTA can be extended to parent-child trios to detect the paternal genetic effect (genetic nurturing effect)^20^. In combination with existing approaches such as M-GCTA and Trio-GCTA^21,24,27^, H-GCTA can also be used to resolve the contribution of POEs and correlations between maternal and fetal genetic effects. We believe this approach represents a significant enhance to the genetic analytic toolbox of pregnancy-related outcomes that others will also employ moving forward.

## Online Methods

### Datasets and quality control

We used genome wide single nucleotide polymorphism (SNP) data from 10,375 mother-child pairs from five European cohorts to distinguish the maternal-fetal genetic contribution to the phenotypic variance of pregnancy-related outcomes such as gestational duration and fetal size measurements at birth (birth weight, birth length and head circumference) (Supplementary Text, Supplementary Figure 1). The study cohorts included Avon Longitudinal Study of Parents and Children (ALSPAC)^49,50^ from UK, Hyperglycemia and Adverse Pregnancy Outcome study (HAPO)^51^ from UK, Canada, and Australia, Finnish dataset (FIN)^31,52^, Danish Birth Cohort (DNBC)^53^, Norwegian Mother, Father and Child Cohort study (MoBa)^54^ (Supplementary Text, Supplementary Figure 2 and Supplementary Tables 1-4). A detailed description of data sets can be found in Supplementary Text.

Genotyping of DNA extracted from whole blood or swab samples was done on various SNP array platforms such as Affymetrix 6.0, Illumina Human550-Quad, Illumina Human610-Quad, Illumina Human 660W-Quad. SNP array data was filtered based on SNP and sample quality. Quality Control (QC) of genotypes data was performed at two levels – marker level and individual level. Marker level QC was conducted using PLINK 1.9^55^ on the basis of SNP call rate, minor allele frequency (MAF), Hardy-Weinberg Equilibrium (HWE) and individual level QC was done on the basis of call rate per individual, average heterozygosity per individual, sex assignment, inbreeding coefficient. Non-European samples were removed from the study by principal components analysis (PCA) anchored with 1,000 genome samples. Following QC, genotype data of mother-child pairs were phased using SHAPEIT 2^56^. SHAPEIT 2 automatically recognizes pedigree information provided in the input files. When phasing mother/child duos together, the first allele in child was always the transmitted allele from mother and the second one from father. We imputed the pre-phased genotypes for missing genotypes on Sanger Imputation Server using Positional Burrows-Wheeler Transform (PBWT) software^57^. Haplotype reference consortium (HRC) panel was utilized as reference data for imputation purpose^58^. The phasing and mother-child allele transmission of the imputed alleles were retained from the pre-phasing stage.

QC of phenotype data was conducted considering gestational duration as the primary outcome. Pregnancies involving history of risk factors for preterm birth or any medical complication during pregnancy influencing preterm birth, C-sections and non-spontaneous births were excluded. We also excluded, non-singlet pregnancies, pregnancies who self-reported non-European ancestry and children who could not survive > 1 year. Additionally, gestational duration was adjusted for fetal sex; fetal size measurements at birth such as birth weight, birth length and head circumference were adjusted for gestational duration up to third orthogonal polynomial component. Details of genotype and phenotype QC is provided in the Supplementary Text.

### Statistical Method

We used a linear mixed model (LMM) to estimate the SNP-heritability (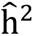) of simulated and empirical phenotypes. This model assumes that the phenotype was normally distributed – Y ∼ N(μ, V) with mean μ and variance V. We created GRMs from standardized genotypes/haplotypes utilizing the method developed by Yang et.al.^7,8^ and Speed et. al.^9,16^. Each cell of the genotype-based GRM and haplotype-based GRM represented relatedness between two individuals j and k calculated based on genotypes (Equation 1) and haplotypes (Equation 2) respectively.

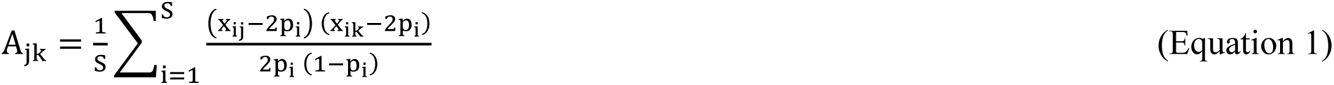

Where, A_jk_ is the correlation coefficient between two individuals j and k averaged over all SNPs; S is number of SNPs used to calculate relatedness; x_ij_ is the number of copies of the reference alleles in individual j for SNP i (i.e. 0 or 1 or 2); x_ik_ is the number of copies of the reference alleles in individual k for SNP i (0 or 1 or 2); p_i_ is frequency of reference allele of SNP i.

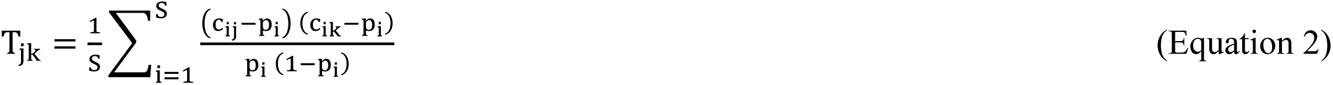

Where, T_jk_ is the correlation coefficient between two mother/child duos or full trios j and k based on maternal transmitted alleles (m1) or maternal non-transmitted alleles (m2) or paternal transmitted alleles (p1) or paternal non-transmitted alleles (p2); S is number of SNPs whose alleles are used to calculate relatedness; c_ij_ is the number of the reference alleles of m1 or m2 or p1 or p2 in mother/child duo or full trio j for SNP i (i.e. 0 or 1); c_ik_ is the number of the reference alleles of m1 or m2 or p1 or p2 in mother/child duo or full trio k for SNP i (i.e. 0 or 1); p_i_ is frequency of reference allele of SNP i.

For genotype-based analysis, we created two GRMs – M and F by utilizing maternal genotypes (m) and fetal genotypes (f) respectively. For haplotype-based analysis, we considered mother-child pair as a single analytical unit consisting of three haplotypes corresponding to m1, m2, and p1. We created three separate GRMs – M1, M2 and P1 using only m1, only m2 and only p1 respectively (Figure 1a, b). We fitted mothers’ genotype-based GRM (M) (Equations 3 and 4) and children’s genotype-based GRM (F) (Equations 5 and 6) separately in LMM to estimate phenotypic variance attributable to maternal and fetal genotypes respectively. To calculate explicit contribution of maternal and fetal genomes to the overall narrow-sense heritability, we simultaneously fitted all three matrices (M1, M2 and P1) in LMM and estimated the additive genetic variance attributable to each of the three components (Equation 7, 8).

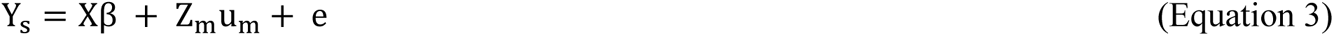

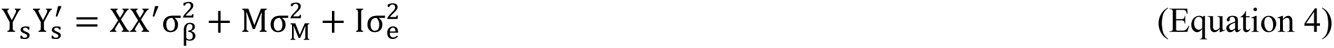

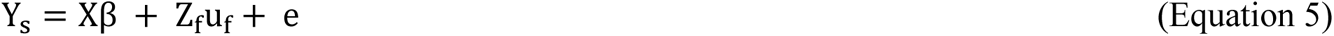

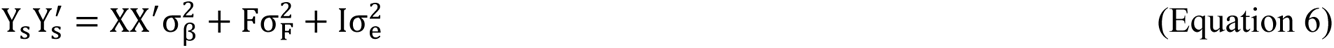

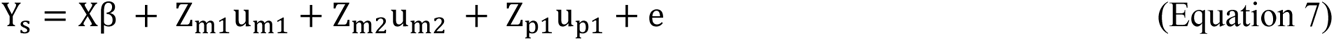

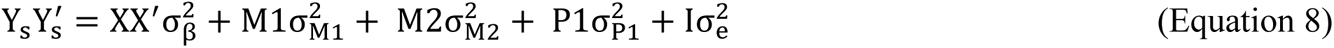

Where, Y_s_ is a vector of standardized phenotype (n x 1; where, n is number of individuals); X is a matrix of covariates representing fixed effects (n x p; where, p is number of fixed effects); β is a vector of fixed effects (p x 1); Z_m_ is a matrix of mothers’ standardized genotypes (m) (n x S; where, S is number of SNPs); Z_f_ is a matrix of children’s standardized genotypes (f) (n x S); Z_m1_ is a matrix of standardized maternal transmitted alleles (m1) (n x S); Z_m2_ is a matrix of standardized maternal non-transmitted alleles (m2) (n x S); Z_p1_ is a matrix of standardized paternal transmitted alleles (p1) (n x S); ε is a vector of residual effects with 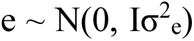; u_m_ and u_f_ are vectors of random effect sizes for maternal genotypes (m) and fetal genotypes (f); u_m1_, u_m2_ and u_p1_ are vectors of random effect sizes for maternal transmitted (m1), maternal non-transmitted (m2) and paternal transmitted (p1) alleles respectively (m x 1); Y Y^′^ is Variance-Covariance matrix of phenotypes; M, F, M1, M2 and P1 are GRMs generated from Z_m_, Z_f_, Z_m1_, Z_m2_ and Z_p1_ respectively 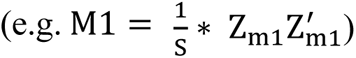; σ^2^ are the variances of the respective components.

As previously reported^9,16,47^, genetic architecture is parametrized on MAF and pair-wise LD, assuming E[var(u_i_)] ∼ w_i_[p_i_(1 − p_i_)]^1+α^, where u_i_, w_i_ and p_i_are the effect size, weight and reference allele frequency of SNP i and α is the scaling factor which represents the extent to which MAF influences the variance of per-allele effect of SNP i [var(u_i_)]. We calculated SNP-specific weights using LDAK and scaled GRMs with two α values (α = −0.25 and − 1.0) in each model. Each standardized column of genotype/haplotype matrix (n x S) was multiplied by w_i_[p_i_ (1 − p_i_)]^1+α^ (α = −0.25, −1.0) before fitting into LMM.

### Implementation

Phenotypic variance i.e. Var(Y) attributable to different components could be estimated by fitting GRMs corresponding to those components in LMM. We used REML implemented through GCTA^7,8^ and LDAK^9,16^ to estimate 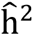 of simulated and empirical phenotypes. For genotype-based analysis through conventional GCTA approach^7^, we fitted a GRM generated from mothers’ genotypes (M) and children’s genotypes (F) separately in LMM whereas for haplotype-based analysis through H-GCTA approach, we fitted three GRMs (M1, M2 and P1) simultaneously in LMM. We also compared results from our approach with those from a contemporary approach, M-GCTA^21,24^. Analysis through the M-GCTA approach involved generation of the GRMs using mothers’ and children’s genotypes together. The upper left quadrant of the GRM represented genetic relationship matrix of mothers (M’); the lower right quadrant represented genetic relationship matrix of children (G) and sum of the lower left quadrant and its transpose represented the genetic relationship matrix of mothers and children (D).

Each approach was fitted through three different models, namely, GREML, LDAK-Thin (where, all pruned SNPs with r^2^ ≤ 0.98 were given equal weights i.e. 1.0) and LDAK-Weights (where, specific weights were calculated for each pruned SNP based on its pair-wise LD with other SNPs in a 100 kb window) (Supplementary Figure 1). For GREML and LDAK-Thin model, constant values of w_i_were used (w_i_ = 1). The difference between GREML and LDAK-Thin model exists in the number of SNPs used to calculate GRM. While GREML uses all genotyped/imputed SNPs to calculate GRM, LDAK-Thin uses only pruned SNPs for the same. On the other hand, LDAK-Weights model uses SNP-specific weights along with specific values of α for scaling.

### Simulation

A total of 100 replicates of phenotypes were simulated using empirical genotype data from 10,375 mother-child pairs. We randomly selected 10,000 causal variants from a common set of all polymorphic SNPs across all datasets (approximately 11 million markers) and randomly picked their effect sizes from standard normal distribution [N(0,1)]. Phenotypes were generated from the model y_j_ = g_j_ + e_j_, where y_j_, g_j_and e_j_ are phenotypic, genetic and residual (environmental) values for individual j. Genetic value of individual j was calculated as 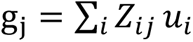 where *Z*_*ij*_ is standardized genotypic value and *u*_*i*_ is effect size of variant i in individual j. Multiplication of randomly picked effect sizes [(*u*_*i*_∼(0,1)] with standardized genotype/haplotype matrix implies that effect sizes are inversely proportional to MAF. A total of 100 independently generated residual values were added to individual’s genetic value (g_j_) to simulate 100 replicates of phenotype. Residual effects were randomly drawn from a distribution 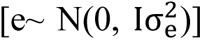 where e is a vector of residual effects, I is an identity matrix and 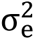 is the variance of residual effects with 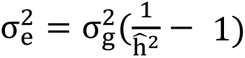 where 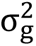 is the variance of genetic values and 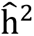 is a preset SNP-based narrow-sense heritability 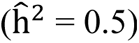.

Three types of traits were simulated considering effects only from the mother (maternal traits), only from fetus (fetal traits) and joint maternal-fetal effects (Table 1). Traits with joint maternal-fetal effects were simulated with different levels of average correlation among maternal and fetal genetic effects (–0.5, –1.0, 0.5 and 1.0) (Table 1). First, traits with independent maternal-fetal genetic effects were simulated using independent and same set sets of causal variants in mothers and children. As we observed similar results in both scenarios, traits with correlated maternal-fetal genetic effects were simulated using same set of 10,000 causal variants in mothers and children. We also simulated fetal traits with POEs, where m1 had less effect in comparison to p1. We considered different scenarios, where varying fractions of causal variants e.g. 25%, 50%, showed maternal imprinting. In each scenario, we simulated different levels of imprinting for m1 (25% – 100%) by reducing effect sizes of m1 (75% – 0%) as compared to p1 (Table 1). Non-zero effects of m1 as compared to p1 represented partial maternal imprinting whereas no effect of m1 represented complete imprinting. All relatedness matrices using simulated data were generated and fitted using different models such as GREML, LDAK-Thin and LDAK-weights into LMM in a similar way as mentioned in the statistical method and Implementation section. We compared our haplotype-based approach (H-GCTA) with conventional GCTA approach and a contemporary M-GCTA approach using above mentioned models with two α values (–1.0, –0.25) for all simulated traits. All analyses for simulated data were run using unrestricted REML i.e. 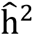 estimates could be less than zero.

### Analysis of Empirical datasets

We performed analyses using three sets of markers – all polymorphic SNPs, SNPs with MAF > 0.001 and SNPs with MAF > 0.01, to include the contribution of very rare, rare and common variants to the heritability of pregnancy-related outcomes (Supplementary Figure 1). The marker sets based on the MAF cutoff were selected in each dataset separately, considering mothers as founders. Then, a common set of markers across all datasets was selected in each MAF cutoff category. We pooled individual datasets and generated five different GRMs utilizing mothers’ genotypes (M), children’s genotypes (F), maternal transmitted haplotypes (M1), maternal non-transmitted haplotypes (M2) and paternal transmitted haplotypes (P1) using the imputed genotype data of mother/child pairs (Supplementary Table 2). One of the related individuals was removed from each GRM (relatedness coefficient > 0.05) and a common set of mother-child pairs across five GRMs was selected in each MAF cutoff category (Supplementary Table 3). The GRMs were created and fitted into LMM using GREML, LDAK-Thin and LDAK-Weights model. LMM-based analyses for empirical data were performed using restricted REML i.e. 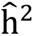 estimates could not be less than zero. All the analyses were adjusted for principal components (PCs) – 20 PCs for analyses through GCTA and M-GCTA and 30 PCs (10 PCs corresponding to m1, m2 and p1 each) for analyses through H-GCTA (Supplementary Figure 6). We also replicated our findings in another Nordic dataset (HARVEST) of ∼ 8,000 mother-child pairs (Supplementary Text). We estimated the 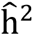 of gestational duration through GREML (α = –1.0) in replication dataset using SNPs with MAF > 0.01 (Supplementary Figure 7, Supplementary Table 25).

## Acknowledgements

We are extremely grateful to all the families who participated in Avon Longitudinal Study of Parents And Children (ALSPAC), Hyperglycemia and Adverse Pregnancy Outcome study (HAPO), Finnish Birth Cohort (FIN), Danish Birth Cohort (DNBC) and Norwegian Mother, Father and Child Cohort study (MoBa), the clinical staff for their consistent help, the whole team of respective studies including interviewers, computer and laboratory technicians, clerical workers, research scientists, volunteers, managers, receptionists and nurses. We also thank organizing bodies for administrating the studies. Our sincere thanks to dbGaP for depositing and hosting data access for the current research.

## Author Contributions

*A. K. S.* contributed in concept and design of the study, analysis and interpretation of data and drafted the work.

*J. J.* contributed in analysis and interpretation of data and substantially revised the manuscript.

*P. S. N.* contributed in interpretation of data and substantially revised the manuscript.

*J.C.* contributed in interpretation of data and substantially revised the manuscript.

*J.B.* contributed in acquisition and analysis of genotype and phenotype data.

*K.T.* contributed in acquisition and analysis of genotype and phenotype data.

*M.H.* contributed in acquisition of genotype and phenotype data and substantially revised the manuscript.

*P. R. N.* contributed in acquisition and analysis of genotype and phenotype data.

*D. M. E.* contributed in interpretation of data and substantially revised the manuscript.

*B.J.* contributed in acquisition of genotype and phenotype data and substantially revised the manuscript.

*L. J. M.* contributed in acquisition of genotype and phenotype data, analysis and interpretation of data and substantially revised the manuscript.

*G. Z.* contributed in concept and design of the study, acquisition of genotype and phenotype data, analysis and interpretation of data and substantially revised the manuscript.

