## Supplementary Text & Figures for "Haplotype-based analysis distinguishes maternal-fetal genetic contribution to pregnancy-related outcomes"

1) Division of Human Genetics, Cincinnati Children's Hospital Medical Center, USA; The Center for Prevention of Preterm Birth, Perinatal Institute, Cincinnati Children's Hospital Medical Center, USA; March of Dimes Prematurity Research Center Ohio Collaborative, USA; Department of Pediatrics, University of Cincinnati College of Medicine, USA; 2) Department of Obstetrics and Gynecology, Institute of Clinical Sciences, Sahlgrenska Academy, University of Gothenburg, Gothenburg, Sweden; 3) Division of Biomedical Informatics, Cincinnati Children's Hospital Medical Center, Cincinnati, OH, USA; 4) Region Västra Götaland, Sahlgrenska University Hospital, Department of Obstetrics and Gynecology, Gothenburg, Sweden; 5) Obstetrics and Gynecology, University of Helsinki and Helsinki University Hospital, Helsinki, Finland; 6) PEDEGO Research Unit and Medical Research Center Oulu, University of Oulu and Department of Children and Adolescents, Oulu University Hospital, Oulu, Finland; 7) KG Jebsen Center for Diabetes Research, Department of Clinical Science, University of Bergen, Bergen, Norway; Division of Health Data and Digitalization, Department of Genetics and Bioinformatics, Norwegian Institute of Public Health, Oslo, Norway; Center for Medical Genetics and Molecular Medicine, Haukeland University Hospital, Bergen, Norway; 8) Institute for Molecular Bioscience, The University of Queensland, Brisbane, Australia; Frazer Institute, The University of Queensland, Brisbane, Australia.; Medical Research Council Integrative Epidemiology Unit, University of Bristol, UK; Bristol Medical School, Population Health Sciences, University of Bristol, Bristol, UK.; 9) Department of Obstetrics and Gynecology, Sahlgrenska Academy, University of Gothenburg, Gothenburg, Sweden.; 10) Department of Genetics and Bioinformatics, Area of Health Data and Digitalization, Norwegian Institute of Public Health, Oslo, Norway

ix) Supplementary Table 9: SNP-based heritability of simulated traits from ALSPAC dataset with correlated maternal-fetal genetic effects (average correlation = -1.0) 16
x) Supplementary Table 10: SNP-based heritability of simulated traits from ALSPAC dataset with correlated maternal-fetal genetic effects (average correlation = -0.5) 17
xi) Supplementary Table 11: SNP-based heritability of simulated traits from ALSPAC dataset with correlated maternal-fetal genetic effects (average correlation = 1.0) 17
xii) Supplementary Table 12: SNP-based heritability of simulated traits from ALSPAC dataset with correlated maternal-fetal genetic effects (average correlation = 0.5) 17
xvii) Supplementary Table 17: SNP-based heritability of simulated traits from pooled dataset with correlated maternal-fetal genetic effects (average correlation = -1.0) 19
xviii) Supplementary Table 18: SNP-based heritability of simulated traits from pooled dataset with correlated maternal-fetal genetic effects (average correlation = -0.5) 19
xix) Supplementary Table 19: SNP-based heritability of simulated traits from pooled dataset with correlated maternal-fetal genetic effects (average correlation = 1.0) 20
xx) Supplementary Table 20: SNP-based heritability of simulated traits from pooled dataset with correlated maternal-fetal genetic effects (average correlation = 0.5) 20
xxv) Supplementary Table 25: Replication of heritability estimation of gestational duration 22

### A) SUPPLEMENTARY TEXT

#### i) Description of datasets

##### a) Avon Longitudinal Study of Parents and Children (ALSPAC)

ALSPAC is an observational study to investigate the genetic and environmental influence on health and development of parents and their offspring[1,2]. The study was approved by ALSPAC Ethics and law Committee and the local Research Ethics Committees. The study consists of 14,541 pregnancies recruited from Avon County, Bristol, England with expected dates of delivery between April 01' 1991 and December 31' 1992. Of these, initial pregnancies, there was a total of 14,676 fetuses, resulting into 14,062 live births and 13,988 children who lived more than 1 year. The women, their partners and children were followed up for 19-22 years with 20 completed questionnaires. The study website contains details of all the data that is available through a fully searchable data dictionary and variable search tool (http://www.bristol.ac.uk/alspac/researchers/our-data/). Genotype data of the mothers and children were generated using the Illumina HumanHap550 quad (children) and Illumina human660W quad (mothers). Genotype data consisted of 17,842 participants (either mothers or children), containing 6,305 mother-child pairs, each with 465,740 SNPs genotyped. A total of 5,369 mother-child pairs who passed genotype QC and inclusion/exclusion criteria were included in the analysis (Supplementary Table 1; Supplementary Figure 2a).

##### b) Hyperglycemia and Adverse Pregnancy Outcome (HAPO)

HAPO is an international study conducted at multiple centers for fetal growth and maternal glucose levels during pregnancy[3]. High quality phenotypic data has been collected from 25,000 pregnant women using standardized protocols, uniform across centers. These women belonged to different racial and socio-demographic backgrounds. We downloaded data of 1,500 mother-child pairs (2,866 samples with genotype data) of European ancestry (phs000096.v2.p1) from dbGaP (http://www.ncbi.nlm.nih.gov/sites/entrez?db=gap/) for the current study. Gestational duration in this dataset was determined by last menstrual period (LMP) or ultrasound estimation between 6-24 weeks. After quality control of genotypes and phenotypes, we included 1089 mother-child pairs for further analysis (Supplementary Table 1; Supplementary Figure 2b).

##### c) Finnish Dataset (FIN)

FIN is a case-control study of spontaneous preterm birth. More than 1,600 mother/child pairs from the Helsinki University Hospitals were recruited between 2004 and 2014[4,5]. The study was approved by the Ethics Committee of Oulu University Hospital and that of Helsinki University Central Hospital. Whole blood samples were collected from all of them after written informed consent and 2,962 blood samples from mothers and children were genotyped. All the studied samples were of Finnish descent. Gestational duration was determined by crown-rump length (CRL) at the first ultrasound screening between 10 and 13 weeks. 1,169 mother-child pairs were selected for further analysis after genotype quality control (QC) procedure and applying the phenotype-based inclusion/exclusion criteria (Supplementary Table 1; Supplementary Figure 2c).

##### d) Danish National Birth Cohort (DNBC)

DNBC is an epidemiological birth cohort for the study of health outcomes in mothers and their offspring[6]. The study followed more than 100,000 pregnancies between 1996 and 2003, starting in the first trimester of pregnancy. The study was approved by the Danish Scientific Ethical Committee and the Danish Data Protection Agency. We used genotype data (phs000103.v1.p1) generated in two genome-wide association studies - the study of preterm delivery and the study of obesity, available at dbGaP website (http://www.ncbi.nlm.nih.gov/sites/entrez?db=gap/). In total, 5,921 mothers and 2,130 infants (1739 mother-child pairs) after QC were utilized for the analysis (Supplementary Table 1; Supplementary Figure 2d). Gestational duration in this dataset was determined by a consensus algorithm combining all available information from multiple sources: self-reported date of last menstrual period (LMP), self-reported delivery date, and gestational duration at birth registered in the Medical Birth Register and the National Patient Register.

##### e) Norwegian Mother and Child Cohort Study (MoBa)

MoBa is a Norwegian pregnancy study for causes of disease in mothers and their offspring, initiated and followed by the Norwegian Institute of Public Health (NIPH)[7]. The study includes more than 114,000 children, 95,000 mothers and 75,000 fathers recruited between 1999 through 2008 starting from week 17 in pregnancy. The study was approved by The Regional Committee for Medical and Health Research Ethics in South-Eastern, Norway (2009/1387 and 2010/2683 S-6075). An informed written consent was obtained prior to sample collection. For most pregnancies, gestational duration was estimated by anatomical ultrasound at gestational weeks 17–19. However, ultrasound dating could not be obtained for some pregnancies and gestational duration was estimated using LMP. For the current study, we used the mother-child pairs, selected from Version 4 of the MoBa dataset, which included a total of 71,669 pregnancies. We selected singleton live-births from mothers in the age group 20–34 years. Pregnancies involving pre-existing medical conditions, complications during pregnancies and conceived by in-vitro fertilization were excluded from the study. Random sampling was done from two gestational duration ranges 154-258 days (cases) and 273–286 days (controls). In total, 3,121 mothers and children were genotyped. 1,834 mothers and 1,143 children (1080 mother-child pairs) were included in the analysis following genotype QC and phenotype inclusion/exclusion (Supplementary Table 1; Supplementary Figure 2e).

#### ii) Quality control of genotypes

Genotyping was performed on various Affymetrix and Illumina SNP array platforms. Specifically, genotyping in ALSPAC dataset was conducted on Illumina HumanHap550 quad (children) and Illumina human660W quad (mothers); SNPs in MoBa dataset were genotyped using Illumina Human660W-Quad bead arrays; and SNPs genotyping of the Finnish cohort was done using Affymetrix SNP array 6.0 and different Illumina arrays. HAPO and DNBC datasets were downloaded from dbGaP. HAPO dataset was genotyped on Human610-Quad array; and DNBC samples were genotyped using Illumina Human660W-Quad bead arrays. Genotype calls from Affymetrix SNP array 6.0 were determined using CRLMM algorithm among chips that passed the vendor-suggested QC (contrast QC > 0.4). Similar procedure was used for Illumina SNP arrays in all datasets except FIN where genotype calling was conducted using Illumina's genotyping module v1.94 in the GenomeStudio v2011.1.

Following genotype calls, we performed genotype QC across all the datasets separately. Genotype QC was performed at individual and marker level using Plink 1.9[8]. Individual-level QC was based on call rate per individual - samples with < 95% SNPs called were excluded from the study, average heterozygosity across all genotypes - samples with substantially high or low heterozygosity were excluded from the study, sex assignment using heterozygosity in X-chromosome SNPs and IBD analysis. Homogeneity of samples and genetic ancestry was determined by principal components analysis (PCA) anchored by 1000 Genomes reference samples. Individuals with non-European ancestry were excluded. Marker level QC was performed on the basis of call rate - SNPs with call rate <98% were excluded from the study, minor allele frequency (MAF) – SNPs with MAF < 0.01 were excluded from the study and Hardy-Weinberg Equilibrium (HWE) – SNPs showing significant deviation from HWE ($P < 5\times10^{-6}$) were excluded[9].

The genotype data of the mothers and children who passed QC was phased together using Shapeit2 to infer maternal transmitted alleles (m1), maternal non-transmitted alleles (m2) and paternal transmitted alleles (p1)[10]. While phasing mother-child together, SHAPEIT automatically detects their relationship provided in the input file. The first allele in child is maternal transmitted whereas second allele is paternal transmitted. In case of trios, the first allele in child is paternal transmitted whereas second allele is maternal transmitted. Then, we imputed the pre-phased mother-child data for missing SNPs using haplotype reference consortium (HRC) data containing 64,976 haplotypes with 39,235,157 SNPs[11]. Phased genotype data was uploaded to Sanger Imputation Server where imputation was done using Positional Burrow-Wheeler Transformation (PBWT)[12]. Since genotype data of the FIN dataset was generated using multiple platforms, the pre-phasing and imputation were done separately for each platform. Later, all samples with imputed genotypes were merged and re-phasing of SNPs with MAF > 0.05 was done using Shapeit2. Pre-phased Finnish dataset was imputed in the same way as earlier. From imputed genotypes, three different sets of SNPs were selected based on MAF cutoff (all polymorphic SNPs, SNPs with MAF > 0.001 and SNPs with MAF > 0.01) in individual datasets separately. Mothers were considered as founders and same set of SNPs were selected from children in each dataset. Then, a common set of SNPs were selected across datasets in each MAF cutoff category. Eventually, genotype data of common sets of SNPs from each dataset were pooled together for further analysis (Supplementary Table 2, 3).

##### iii) Quality control of phenotypes

In the current study, we focused on pregnancy outcomes including gestational duration and fetal size measurements at birth (birth weight, birth length and head circumference). All traits in current study were available only in two datasets – ALSPAC and HAPO. Birth weight was accessible in all datasets except MoBa. Birth length was accessible in ALSPAC, HAPO and FIN whereas head circumference was available in the ALSPAC and HAPO. We considered gestational duration as primary trait while doing quality control (QC) of phenotypes. We only included spontaneous, singleton pregnancies whose children were alive > 1 year and both parents self-reported European ancestry. We excluded pregnancies with history of medical conditions influencing pre-term birth such as pre-pregnancy diabetes, hypertension, placental and congenital anomalies. We also excluded pregnancies with any risk factors for pre-term birth during pregnancies such as gestational diabetes, gestational hypertension and preeclampsia (Supplementary Table 4).

**iv) Heritability estimation using SNPs with MAF > 0.001 and MAF > 0.01**

Besides all polymorphic SNPs, $\hat{h}^2$ was also estimated using SNPs with MAF > 0.001 and MAF > 0.01 in pooled dataset of 10,375 unrelated mother-child pairs (relatedness coefficient cutoff > 0.05). The focus of these analyses was to estimate the contribution of very rare, rare and common variants. We estimated $\hat{h}^2$ of gestational duration and gestational duration adjusted birth weight, birth length and head circumference using conventional GCTA[13,14], contemporary M-GCTA[15,16] and newly developed H-GCTA approach by utilizing REML[17,18] implemented through GCTA and LDAK. We used different models accounting for the influence of pair-wise linkage disequilibrium (LD) and minor allele frequency (MAF) on $\hat{h}^2$ estimates (Supplementary Table 23-24). Results obtained from these analyses were like the results obtained through analysis using all polymorphic SNPs. These results indicated that exclusion of very rare and rare variants does not substantially change heritability estimates.

**v) Replication of heritability estimation**

We also replicated analysis using our approach through GREML ($\alpha$ = -1.0) implemented through GCTA in another Nordic cohort – HARVEST from Norway. In this study, approximately 8,000 mother-child pairs were available after quality control. Standard procedures similar to MoBa were used while blood sample collection and genotyping. Only gestational duration was available from the dataset. We used common SNPs with MAF > 0.01 for SNP-based narrow-sense heritability ($\hat{h}^2$) estimation. Similar, to discovery cohorts, gestational duration was adjusted for fetal sex. We observed genetic variance attributable to maternal transmitted alleles (m1) – 7.7% (S.E.: 3.8%, p value = 2.22E-02), maternal non-transmitted alleles (m2) – 4.3% (S.E.: 3.8%, p value = 1.30E-01) and paternal transmitted alleles (P1) – 2.5% (S.E.: 3.7%, p value = 2.49E-02). Results from HARVEST dataset confirmed our findings that genetic variance of gestation duration is mainly contributed by maternal genome (Supplementary Figure 7; Supplementary Table 25).

#### B) Supplementary Figures

##### i) Framework of the study

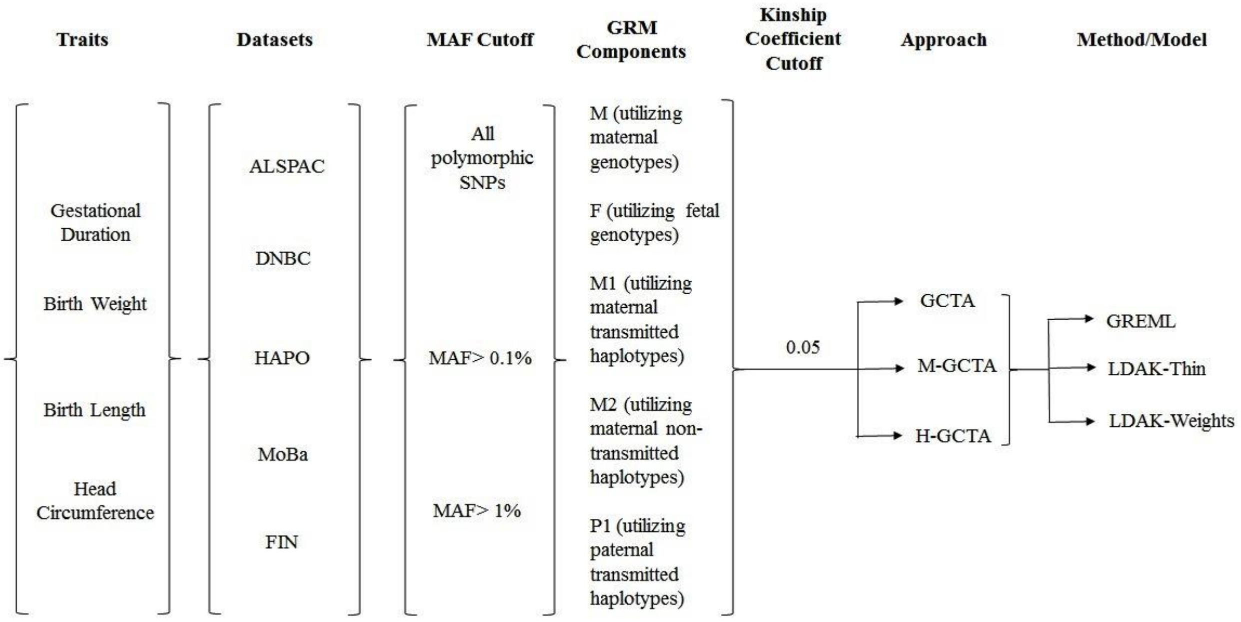

**Supplementary Figure 1:** Framework of the study depicting the traits under study, available datasets, MAF cutoffs, list of GRMs created in each MAF cutoff category, selection of unrelated mother-child pairs. Last block shows methods/models utilized for estimation and comparison of $\hat{h}^2$ estimated from our approach (H-GCTA) with those obtained by two available approaches – GCTA and M-GCTA.

#### ii) Distribution of available phenotypes in datasets

##### a) Phenotypes distribution in ALSPAC dataset

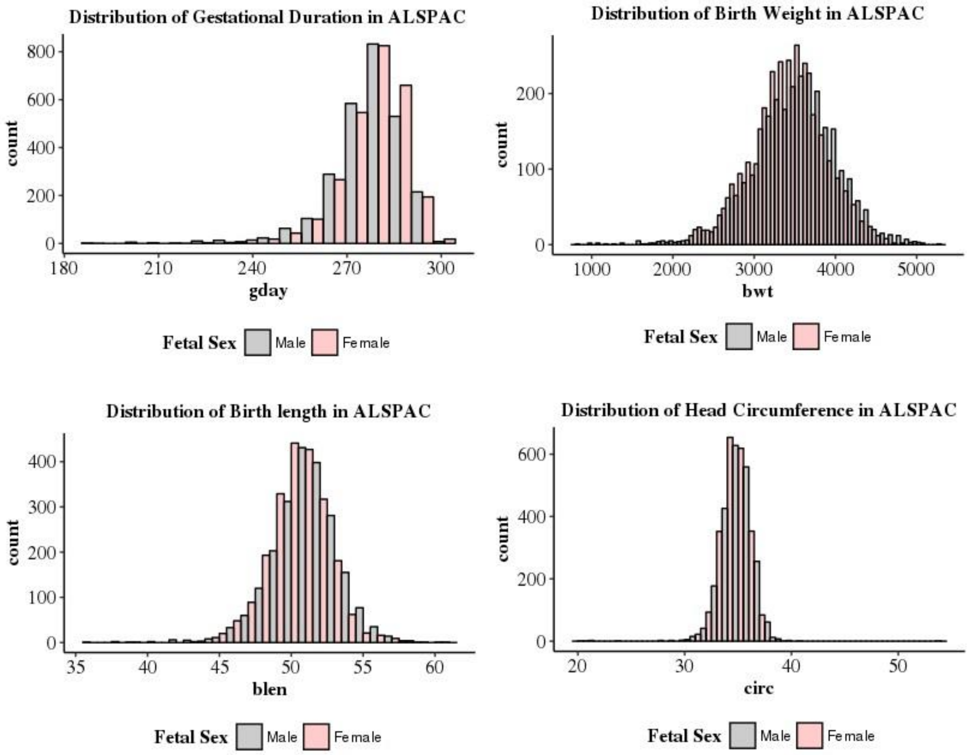

##### b) Phenotypes distribution in HAPO dataset

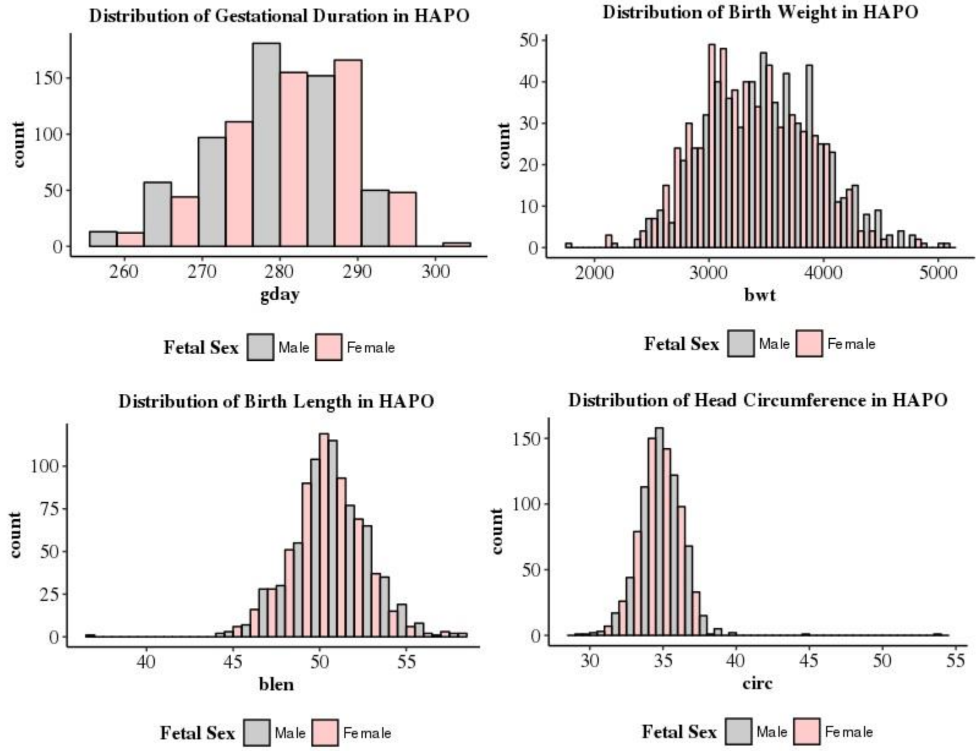

#### c) Phenotypes distribution in FIN dataset

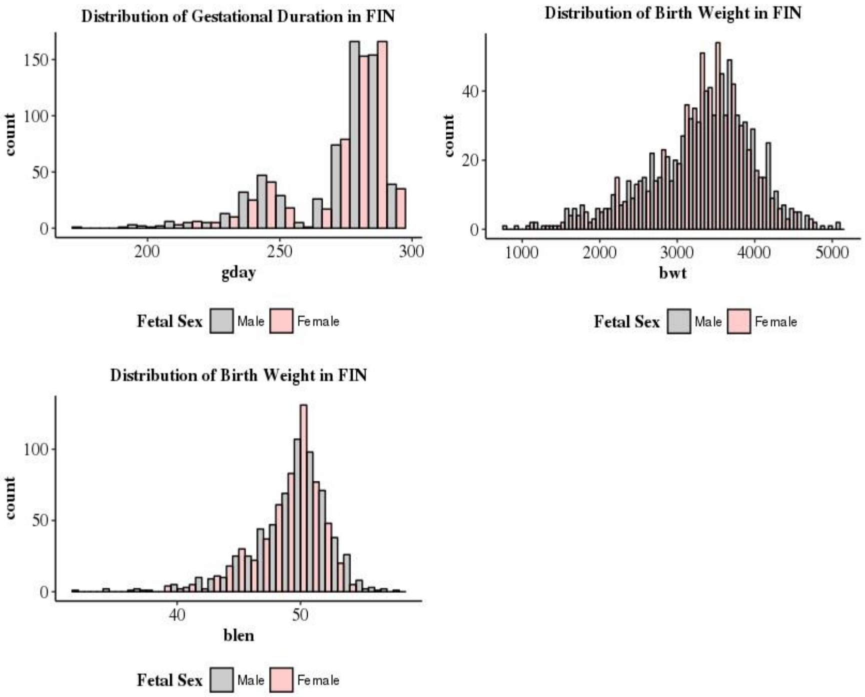

#### d) Phenotypes distribution in DNBC dataset

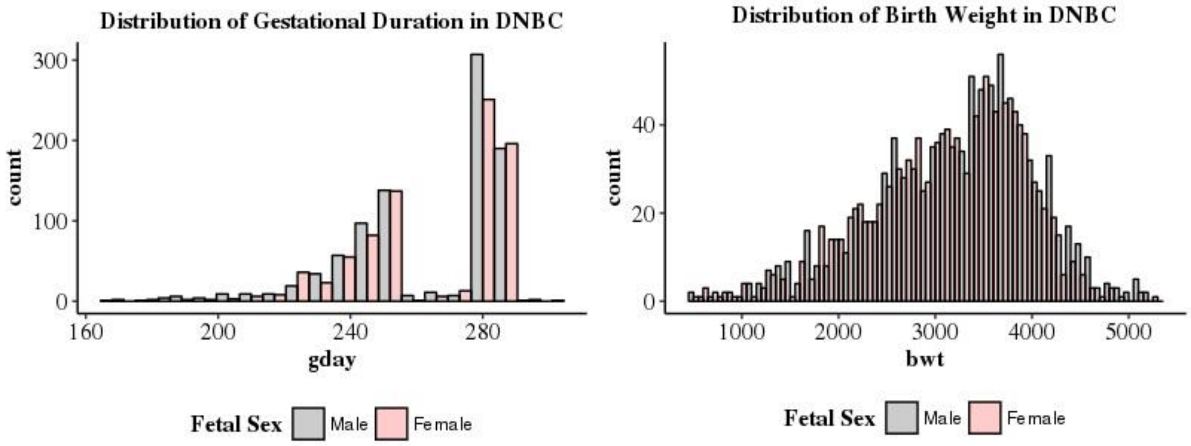

**e) Phenotypes distribution in MoBa dataset**

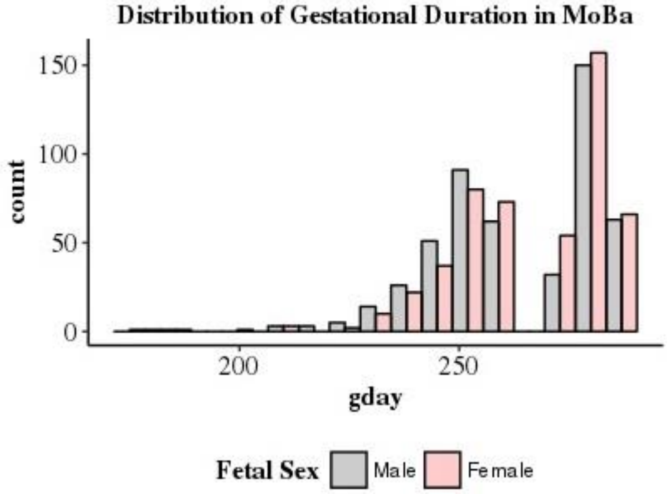

**Supplementary Figure 2:** Distribution of available phenotypes in each dataset categorized by fetal sex – a) distribution of gestational duration, birth weight, birth length and head circumference in ALSPAC dataset; b) distribution of gestational duration, birth weight, birth length and head circumference in HAPO dataset; c) distribution of gestational duration, birth weight and birth length in FIN dataset; d) distribution of gestational duration and birth weight in DNBC dataset and e) distribution of gestational duration in MoBa dataset.

##### iii) Comparison of $\hat{h}^2$ for simulated traits from ALSPAC dataset – maternal traits, fetal traits and traits with independent maternal-fetal genetic effects

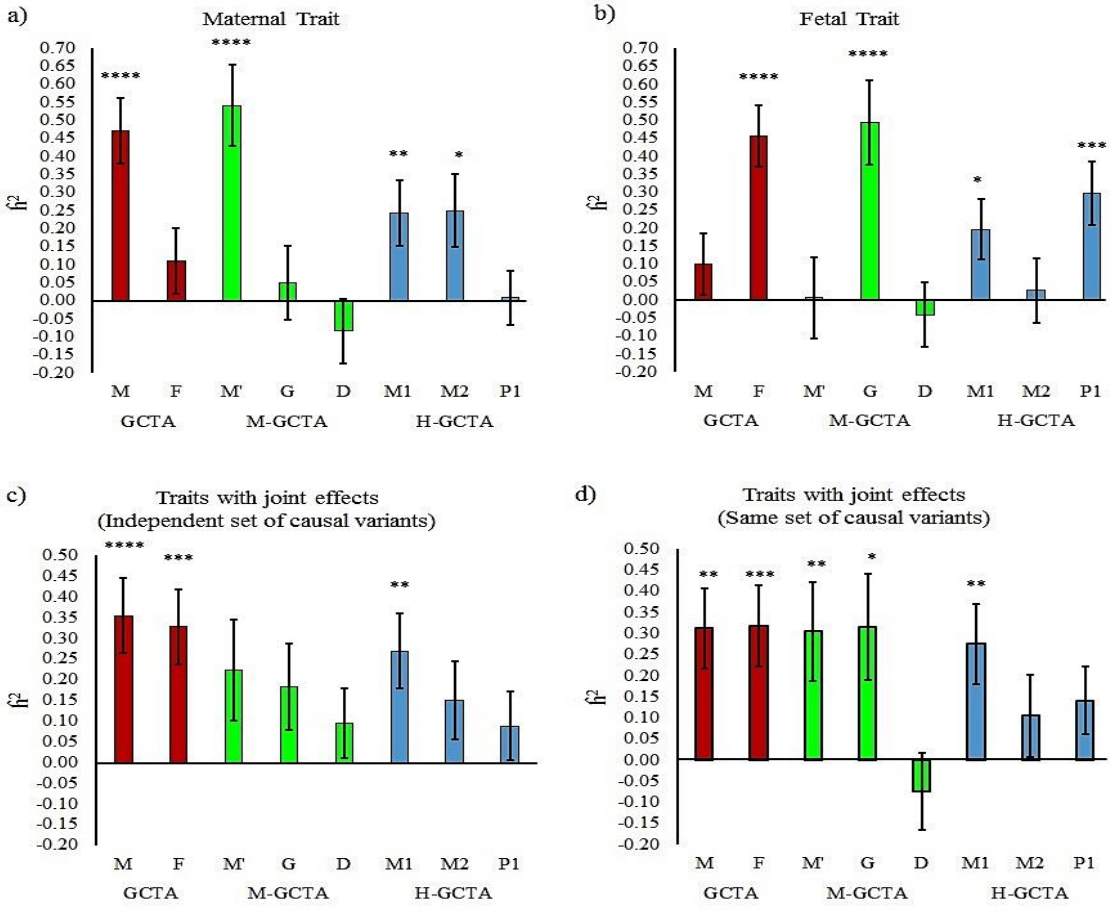

**Supplementary Figure 3:** Comparison of $\hat{h}^2$ for simulated traits from ALSPAC dataset, estimated through different approaches fitting GREML (α = -1.0): a) maternal traits; b) fetal traits; c) traits where independent sets of causal variants have effects through mother and fetus; d) traits where same set of causal variants have effects through mother and fetus. For GCTA, M is the GRM generated from maternal genotypes (m), and F is the GRM generated from fetal genotypes (f). For M-GCTA, M' represents the genetic relationship matrix of mothers; G represents genetic relationship matrix of children and D represents mother-child covariance matrix. For H-GCTA, M1 is the GRM generated from maternal transmitted alleles (m1), M2 is the GRM generated from maternal non-transmitted alleles (m2), and P1 is the GRM generated from paternal transmitted alleles (p1). A total of 100 replicates of each phenotype were simulated using empirical genotypes of ALSPAC dataset. P-values were calculated using z test statistics (two sided). * = (p value < 5.0E-02), ** = (p value < 1.0E-02), *** = (p value < 1.0E-03) and **** = (p value < 1.0E-04).

##### iv) Comparison of $\hat{h}^2$ for simulated traits from ALSPAC dataset –traits with correlated maternal-fetal genetic effects

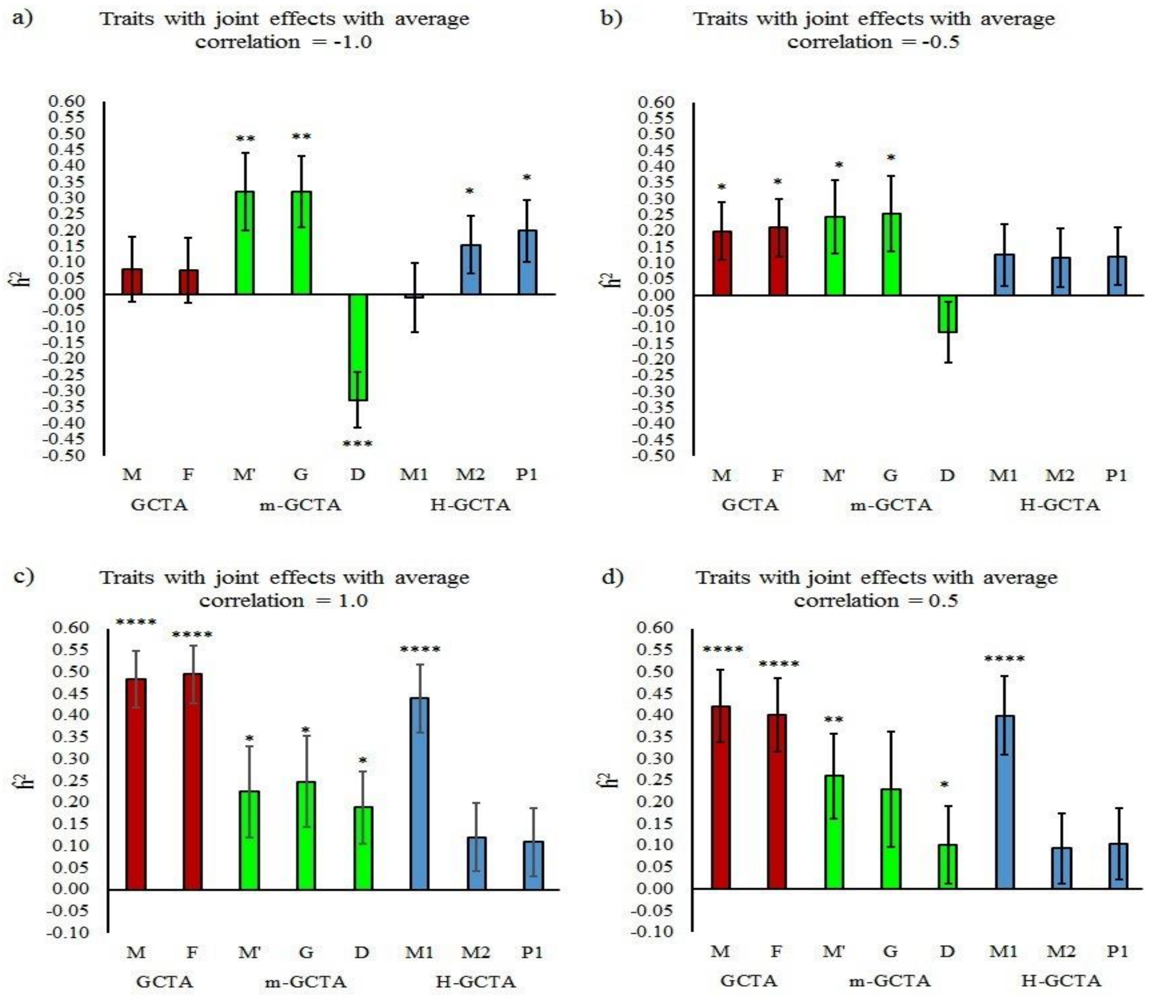

**Supplementary Figure 4:** Comparison of $\hat{h}^2$ estimated through different approaches fitting GREML (α = -1.0) for simulated traits with joint maternal–fetal effects from ALSPAC dataset: a) average correlation = -1.0; b) average correlation = -0.5; c) average correlation = 1.0; d) average correlation = 0.5. For GCTA, M is the GRM generated from maternal genotypes (m), and F is the GRM generated from fetal genotypes (f). For M-GCTA, M' represents the genetic relationship matrix of mothers; G represents genetic relationship matrix of children and D represents mother-child covariance matrix. For H-GCTA, M1 is the GRM generated from maternal transmitted alleles (m1), M2 is the GRM generated from maternal non-transmitted alleles (m2), and P1 is the GRM generated from paternal transmitted alleles (p1). A total of 100 replicates of each phenotype were simulated using empirical genotypes of ALSPAC dataset. P-values were calculated using z test statistics (two sided). * = (p value < 5.0E-02), ** = (p value < 1.0E-02), *** = (p value < 1.0E-03) and **** = (p value < 1.0E-04)

#### v) Schematic representation of variance attributable to maternal transmitted, maternal non-transmitted and paternal transmitted haplotypes in H-GCTA

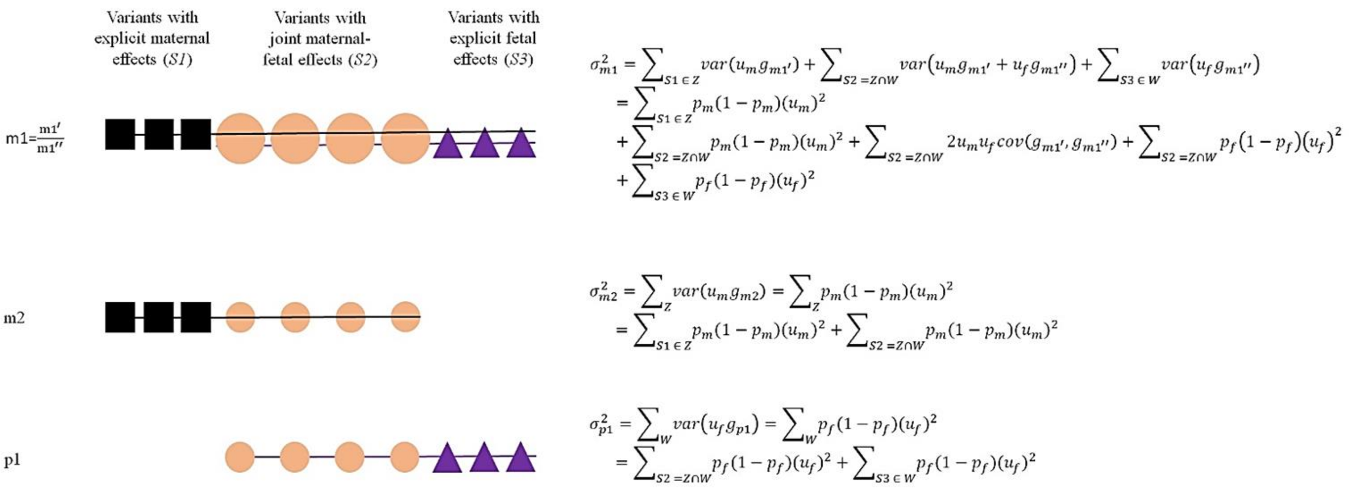

**Supplementary Figure 5:** Schematic representation of variance attributable to maternal transmitted ($\sigma^2_{m1}$), maternal non-transmitted ($\sigma^2_{m2}$) and paternal transmitted ($\sigma^2_{p1}$) haplotypes in H-GCTA – Z and W are the sets of causal variants with maternal and fetal effects, respectively. S1 (Black squares), S2 (orange circles) and S3 (purple triangles) are sets of causal variants with explicit maternal effects, joint-maternal-fetal effects and explicit fetal effects such that S1 ∈ Z, S2 = Z ∩ W and S3 ∈ W. $u_m$ and $u_f$ are causal effects through mother and fetus and $p_m$ and $p_f$ are reference allele frequencies of a causal variant in mother and fetus, respectively. Since each allele is a random draw from Bernoulli distribution, variance in terms of allele frequency is represented as $p_m(1-p_m)$ and $p_f(1-p_f)$ in mother and fetus, respectively. m1 affects the phenotype through maternal transmitted alleles in mother ($g_{m1'}$) and maternal transmitted alleles in fetus ($g_{m1''}$). Likewise, $g_{m2}$ and $g_{p1}$ represent the maternal non-transmitted and paternal transmitted alleles. Therefore, allelic effects - $u_{m1'} = u_{m2}$, $u_{m1''} = u_{p1}$ (in the absence of POEs) and $cov(g_{m1'}, g_{m1''}) = \rho(g_{m1'}, g_{m1''})\sigma_{g_{m1'}}, \sigma_{g_{m1''}}$ is the covariance of two binomial random variables m1' and m1'' present in mother and fetus, respectively; where, ρ and $\sigma$ represent correlation and standard deviation of respective alleles. For a causal variant with joint maternal-fetal effect, $p_m = p_f = p$ in a random mating population, therefore, $cov(g_{m1'}, g_{m1''}) = p(1-p)$ and total phenotypic variance explained by m1, m2 and p1 is $2p(1-p)(u_m^2 + u_f^2 + u_m u_f)$.

##### vi) Principal Components Analysis (PCA) plots using all polymorphic SNPs

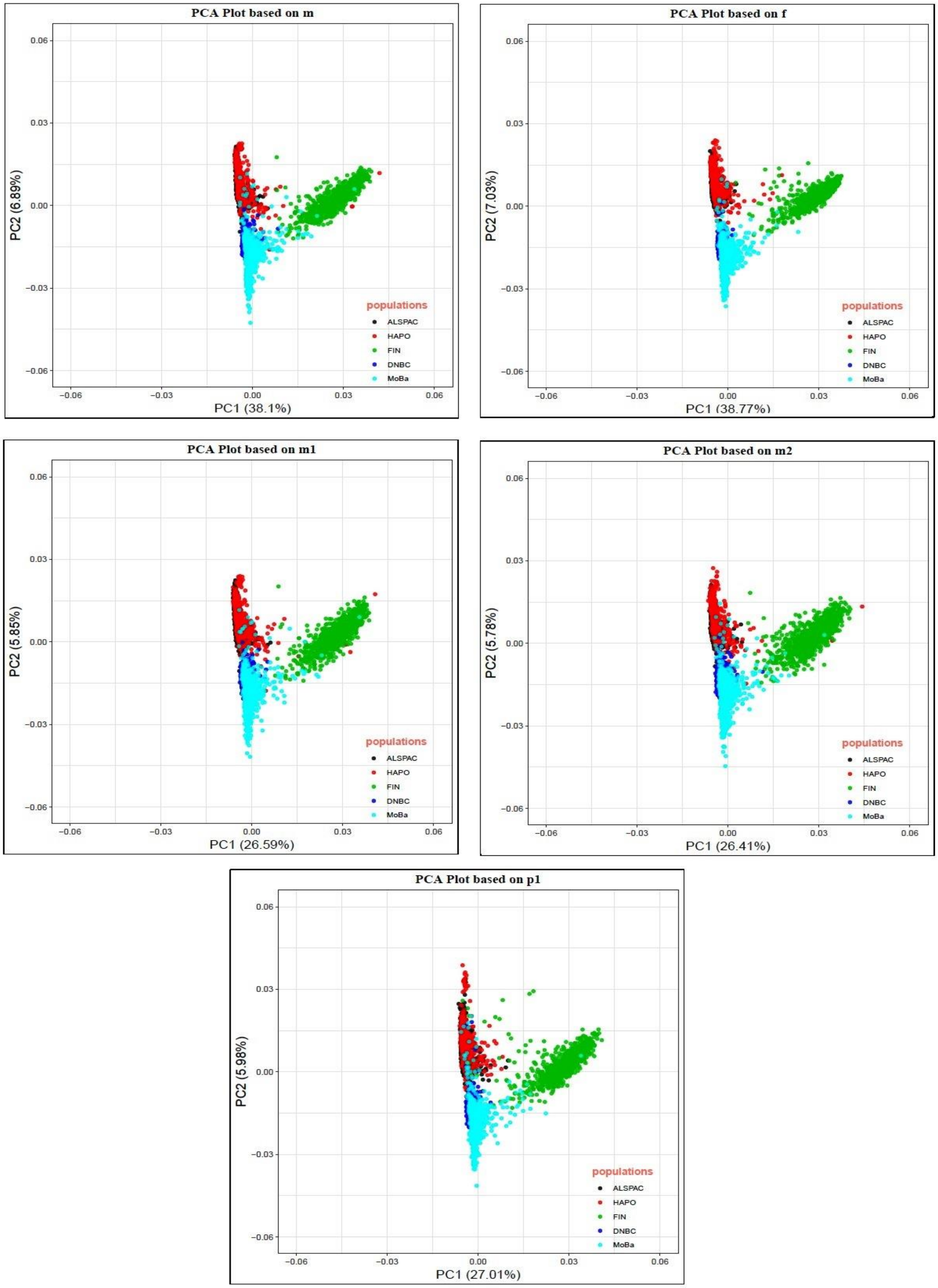

**Supplementary Figure 6:** Principal Components Analysis (PCA) plots of unrelated (relatedness coefficient < 0.5) mother-child pairs using all polymorphic SNPs. m: Mothers' Genotypes; f: children's genotypes; m1: maternal transmitted alleles; m2: maternal non-transmitted alleles; and p1: paternal transmitted alleles.

**vii) Replication of heritability estimation of gestational duration**

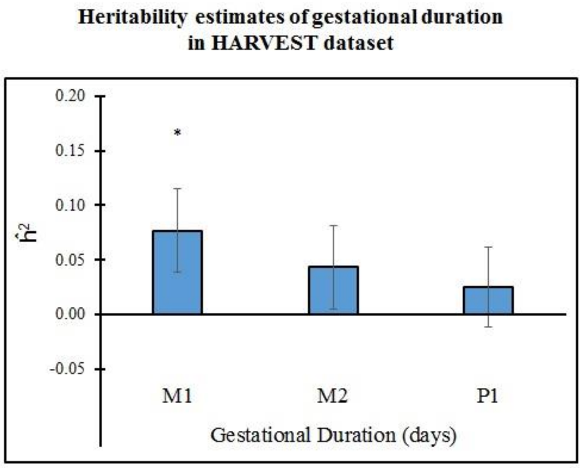

**Supplementary Figure 7:** $\hat{h}^2$ estimation of fetal sex adjusted gestational duration in HARVEST dataset using our approach (H-GCTA). Analysis was performed through GREML (α = -1.0) using SNPs with MAF > 0.01.

#### C) Supplementary tables (Legends)

##### i) Supplementary Table 1: Genotype and Phenotype records in datasets

Number of pregnancies and genotypes present in individual datasets. a) Number of genotypes typed in each dataset b) number of genotypes passed through genotype QC; c) number of pregnancies after genotype QC and phenotype inclusion/exclusion.

##### ii) Supplementary Table 2: Genotype information in individual datasets

Number of imputed sites using Haplotype Reference Consortium (HRC), polymorphic SNPs in individual datasets and common set of SNPs across all available datasets. In individual datasets, mothers were considered as founders for each MAF cutoff category and corresponding children were selected. Final analysis was performed using pooled data and common set of SNPs across all datasets.

##### iii) Supplementary Table 3: Phenotype information in pooled dataset

Number of samples with gestational duration, birth weight, birth length and head circumference in the pooled data; a) without relatedness coefficient cut-off, b) with relatedness coefficient cut-off < 0.05. For mother-child pairs with relatedness coefficient cutoff < 0.05, common set of mother-child pairs were selected from GRMs based on mother's genotypes, children's genotypes, maternal transmitted alleles (m1), maternal non-transmitted alleles (m2) and paternal transmitted alleles (p1).

##### iv) Supplementary Table 4: Phenotype summary in individual datasets

Descriptive statistics of gestational duration, birth weight, birth length and head circumference in ALSPAC, HAPO, FIN, DNBC and MoBa. All four traits were available only in two datasets, namely ALSPAC and HAPO.
Ŧ – Only gestational duration data was available to in MoBa dataset.

##### v) Supplementary Table 5: SNP-based heritability of simulated maternal traits from ALSPAC dataset

$\hat{h}^2$ of simulated maternal traits from ALSPAC dataset, estimated through conventional GCTA, M-GCTA and H-GCTA approach. Each approach was fitted using GREML (α = -0.25, -1.0), LDAK-Thin (α = -0.25, -1.0) and LDAK-Weights (α = -0.25, -1.0). For GCTA, M is the GRM generated from maternal genotypes (m), and F is the GRM generated from fetal genotypes (f). For M-GCTA, M' represents the genetic relationship matrix of mothers; G represents genetic relationship matrix of children and D represents mother-child covariance matrix. For H-GCTA, M1 is the GRM generated from maternal transmitted alleles (m1), M2 is the GRM generated from maternal non-transmitted alleles (m2), and P1 is the GRM generated from paternal transmitted alleles (p1). A total of 100 replicates of each phenotype were simulated using empirical genotypes of ALSPAC dataset. P-values were calculated using z test statistics (two sided).

##### vi) Supplementary Table 6: SNP-based heritability of simulated fetal traits from ALSPAC dataset

$\hat{h}^2$ of simulated fetal traits from ALSPAC dataset, estimated through conventional GCTA, M-GCTA and H-GCTA approach. Each approach was fitted using GREML (α = -0.25, -1.0), LDAK-

Thin (α = -0.25, -1.0) and LDAK-Weights (α = -0.25, -1.0). For GCTA, M is the GRM generated from maternal genotypes (m), and F is the GRM generated from fetal genotypes (f). For M-GCTA, M' represents the genetic relationship matrix of mothers; G represents genetic relationship matrix of children and D represents mother-child covariance matrix. For H-GCTA, M1 is the GRM generated from maternal transmitted alleles (m1), M2 is the GRM generated from maternal non-transmitted alleles (m2), and P1 is the GRM generated from paternal transmitted alleles (p1). A total of 100 replicates of each phenotype were simulated using empirical genotypes of ALSPAC dataset. P-values were calculated using z test statistics (two sided).

##### vii) Supplementary Table 7: SNP-based heritability of simulated traits from ALSPAC dataset with independent maternal-fetal genetic effects using independent sets of causal variants in mother and child

$\hat{h}^2$ of simulated traits from ALSPAC dataset with independent maternal-fetal genetic effects (independent sets of causal variants in mother and child), estimated through conventional GCTA, M-GCTA and H-GCTA approach. Each approach was fitted using GREML (α = -0.25, -1.0), LDAK-Thin (α = -0.25, -1.0) and LDAK-Weights (α = -0.25, -1.0). For GCTA, M is the GRM generated from maternal genotypes (m), and F is the GRM generated from fetal genotypes (f). For M-GCTA, M' represents the genetic relationship matrix of mothers; G represents genetic relationship matrix of children and D represents mother-child covariance matrix. For H-GCTA, M1 is the GRM generated from maternal transmitted alleles (m1), M2 is the GRM generated from maternal non-transmitted alleles (m2), and P1 is the GRM generated from paternal transmitted alleles (p1). A total of 100 replicates of each phenotype were simulated using empirical genotypes of ALSPAC dataset. P-values were calculated using z test statistics (two sided).

##### viii) Supplementary Table 8: SNP-based heritability of simulated traits from ALSPAC dataset with independent maternal-fetal genetic effects using same set of causal variants in mother and child

$\hat{h}^2$ of simulated traits from ALSPAC dataset with independent maternal-fetal genetic effects (same set of causal variants in mother and child), estimated through conventional GCTA, M-GCTA and H-GCTA approach. Each approach was fitted using GREML (α = -0.25, -1.0), LDAK-Thin (α = -0.25, -1.0) and LDAK-Weights (α = -0.25, -1.0). For GCTA, M is the GRM generated from maternal genotypes (m), and F is the GRM generated from fetal genotypes (f). For M-GCTA, M' represents the genetic relationship matrix of mothers; G represents genetic relationship matrix of children and D represents mother-child covariance matrix. For H-GCTA, M1 is the GRM generated from maternal transmitted alleles (m1), M2 is the GRM generated from maternal non-transmitted alleles (m2), and P1 is the GRM generated from paternal transmitted alleles (p1). A total of 100 replicates of each phenotype were simulated using empirical genotypes of ALSPAC dataset. P-values were calculated using z test statistics (two sided).

##### ix) Supplementary Table 9: SNP-based heritability of simulated traits from ALSPAC dataset with correlated maternal-fetal genetic effects (average correlation = -1.0)

$\hat{h}^2$ of simulated traits with correlated maternal-fetal genetic effects (average correlation = -1.0), estimated through conventional GCTA, M-GCTA and H-GCTA approach. Each approach was fitted using GREML (α = -0.25, -1.0), LDAK-Thin (α = -0.25, -1.0) and LDAK-Weights (α = -0.25, -1.0). For GCTA, M is the GRM generated from maternal genotypes (m), and F is the GRM generated from fetal genotypes (f). For M-GCTA, M' represents the genetic relationship matrix of mothers; G represents genetic relationship matrix of children and D represents mother-child covariance matrix. For H-GCTA, M1 is the GRM generated from maternal transmitted alleles (m1), M2 is the GRM generated from maternal non-transmitted alleles (m2), and P1 is the GRM generated from paternal transmitted alleles (p1). A total of 100 replicates of each phenotype were simulated using empirical genotypes of ALSPAC dataset. P-values were calculated using z test statistics (two sided).

**x) Supplementary Table 10: SNP-based heritability of simulated traits from ALSPAC dataset with correlated maternal-fetal genetic effects (average correlation = -0.5)**

$\hat{h}^2$ of simulated traits from ALSPAC dataset with correlated maternal-fetal genetic effects (average correlation = -0.5), estimated through conventional GCTA, M-GCTA and H-GCTA approach. Each approach was fitted using GREML (α = -0.25, -1.0), LDAK-Thin (α = -0.25, -1.0) and LDAK-Weights (α = -0.25, -1.0). For GCTA, M is the GRM generated from maternal genotypes (m), and F is the GRM generated from fetal genotypes (f). For M-GCTA, M' represents the genetic relationship matrix of mothers; G represents genetic relationship matrix of children and D represents mother-child covariance matrix. For H-GCTA, M1 is the GRM generated from maternal transmitted alleles (m1), M2 is the GRM generated from maternal non-transmitted alleles (m2), and P1 is the GRM generated from paternal transmitted alleles (p1). A total of 100 replicates of each phenotype were simulated using empirical genotypes of ALSPAC dataset. P-values were calculated using z test statistics (two sided).

**xi) Supplementary Table 11: SNP-based heritability of simulated traits from ALSPAC dataset with correlated maternal-fetal genetic effects (average correlation = 1.0)**

$\hat{h}^2$ of simulated traits from ALSPAC dataset with correlated maternal-fetal genetic effects (average correlation = 1.0), estimated through conventional GCTA, M-GCTA and H-GCTA approach. Each approach was fitted using GREML (α = -0.25, -1.0), LDAK-Thin (α = -0.25, -1.0) and LDAK-Weights (α = -0.25, -1.0). For GCTA, M is the GRM generated from maternal genotypes (m), and F is the GRM generated from fetal genotypes (f). For M-GCTA, M' represents the genetic relationship matrix of mothers; G represents genetic relationship matrix of children and D represents mother-child covariance matrix. For H-GCTA, M1 is the GRM generated from maternal transmitted alleles (m1), M2 is the GRM generated from maternal non-transmitted alleles (m2), and P1 is the GRM generated from paternal transmitted alleles (p1). A total of 100 replicates of each phenotype were simulated using empirical genotypes of ALSPAC dataset. P-values were calculated using z test statistics (two sided).

**xii) Supplementary Table 12: SNP-based heritability of simulated traits from ALSPAC dataset with correlated maternal-fetal genetic effects (average correlation = 0.5)**

$\hat{h}^2$ of simulated traits from ALSPAC dataset with correlated maternal-fetal genetic effects (average correlation = 0.5), estimated through conventional GCTA, M-GCTA and H-GCTA approach. Each approach was fitted using GREML (α = -0.25, -1.0), LDAK-Thin (α = -0.25, -1.0) and LDAK-

Weights (α = -0.25, -1.0). For GCTA, M is the GRM generated from maternal genotypes (m), and F is the GRM generated from fetal genotypes (f). For M-GCTA, M' represents the genetic relationship matrix of mothers; G represents genetic relationship matrix of children and D represents mother-child covariance matrix. For H-GCTA, M1 is the GRM generated from maternal transmitted alleles (m1), M2 is the GRM generated from maternal non-transmitted alleles (m2), and P1 is the GRM generated from paternal transmitted alleles (p1). A total of 100 replicates of each phenotype were simulated using empirical genotypes of ALSPAC dataset. P-values were calculated using z test statistics (two sided).

##### xiii) Supplementary Table 13: SNP-based heritability of simulated maternal traits from pooled dataset

$\hat{h}^2$ of simulated maternal traits from pooled dataset , estimated through conventional GCTA, M-GCTA and H-GCTA approach. Each approach was fitted using GREML (α = -0.25, -1.0), LDAK-Thin (α = -0.25, -1.0) and LDAK-Weights (α = -0.25, -1.0). For GCTA, M is the GRM generated from maternal genotypes (m), and F is the GRM generated from fetal genotypes (f). For M-GCTA, M' represents the genetic relationship matrix of mothers; G represents genetic relationship matrix of children and D represents mother-child covariance matrix. For H-GCTA, M1 is the GRM generated from maternal transmitted alleles (m1), M2 is the GRM generated from maternal non-transmitted alleles (m2), and P1 is the GRM generated from paternal transmitted alleles (p1). A total of 100 replicates of each phenotype were simulated using empirical genotypes of Pooled dataset. P-values were calculated using z test statistics (two sided).

##### xiv) Supplementary Table 14: SNP-based heritability of simulated fetal traits from pooled dataset

$\hat{h}^2$ of simulated fetal traits from pooled dataset, estimated through conventional GCTA, M-GCTA and H-GCTA approach. Each approach was fitted using GREML (α = -0.25, -1.0), LDAK-Thin (α = -0.25, -1.0) and LDAK-Weights (α = -0.25, -1.0). For GCTA, M is the GRM generated from maternal genotypes (m), and F is the GRM generated from fetal genotypes (f). For M-GCTA, M' represents the genetic relationship matrix of mothers; G represents genetic relationship matrix of children and D represents mother-child covariance matrix. For H-GCTA, M1 is the GRM generated from maternal transmitted alleles (m1), M2 is the GRM generated from maternal non-transmitted alleles (m2), and P1 is the GRM generated from paternal transmitted alleles (p1). A total of 100 replicates of each phenotype were simulated using empirical genotypes of Pooled dataset. P-values were calculated using z test statistics (two sided).

##### xv) Supplementary Table 15: SNP-based heritability of simulated traits from pooled dataset with independent maternal-fetal genetic effects using independent sets of causal variants in mother and child

$\hat{h}^2$ of simulated traits from pooled dataset with independent maternal-fetal genetic effects (independent sets of causal variants in mother and child), estimated through conventional GCTA, M-GCTA and H-GCTA approach. Each approach was fitted using GREML (α = -0.25, -1.0), LDAK-Thin (α = -0.25, -1.0) and LDAK-Weights (α = -0.25, -1.0). For GCTA, M is the GRM generated from maternal genotypes (m), and F is the GRM generated from fetal genotypes (f). For

M-GCTA, M’ represents the genetic relationship matrix of mothers; G represents genetic relationship matrix of children and D represents mother-child covariance matrix. For H-GCTA, M1 is the GRM generated from maternal transmitted alleles (m1), M2 is the GRM generated from maternal non-transmitted alleles (m2), and P1 is the GRM generated from paternal transmitted alleles (p1). A total of 100 replicates of each phenotype were simulated using empirical genotypes of Pooled dataset. P-values were calculated using z test statistics (two sided).

##### xvi) Supplementary Table 16: SNP-based heritability of simulated traits from pooled dataset with independent maternal-fetal genetic effects using same set of causal variants in mother and child

$\hat{h}^2$ of simulated traits from pooled dataset with independent maternal-fetal genetic effects (same set of causal variants in mother and child), estimated through conventional GCTA, M-GCTA and H-GCTA approach. Each approach was fitted using GREML (α = -0.25, -1.0), LDAK-Thin (α = -0.25, -1.0) and LDAK-Weights (α = -0.25, -1.0). For GCTA, M is the GRM generated from maternal genotypes (m), and F is the GRM generated from fetal genotypes (f). For M-GCTA, M’ represents the genetic relationship matrix of mothers; G represents genetic relationship matrix of children and D represents mother-child covariance matrix. For H-GCTA, M1 is the GRM generated from maternal transmitted alleles (m1), M2 is the GRM generated from maternal non-transmitted alleles (m2), and P1 is the GRM generated from paternal transmitted alleles (p1). A total of 100 replicates of each phenotype were simulated using empirical genotypes of Pooled dataset. P-values were calculated using z test statistics (two sided).

##### xvii) Supplementary Table 17: SNP-based heritability of simulated traits from pooled dataset with correlated maternal-fetal genetic effects (average correlation = -1.0)

$\hat{h}^2$ of simulated traits from pooled dataset with correlated maternal-fetal genetic effects (average correlation = -1.0), estimated through conventional GCTA, M-GCTA and H-GCTA approach. Each approach was fitted using GREML (α = -0.25, -1.0), LDAK-Thin (α = -0.25, -1.0) and LDAK-Weights (α = -0.25, -1.0). For GCTA, M is the GRM generated from maternal genotypes (m), and F is the GRM generated from fetal genotypes (f). For M-GCTA, M’ represents the genetic relationship matrix of mothers; G represents genetic relationship matrix of children and D represents mother-child covariance matrix. For H-GCTA, M1 is the GRM generated from maternal transmitted alleles (m1), M2 is the GRM generated from maternal non-transmitted alleles (m2), and P1 is the GRM generated from paternal transmitted alleles (p1). A total of 100 replicates of each phenotype were simulated using empirical genotypes of Pooled dataset. P-values were calculated using z test statistics (two sided).

##### xviii) Supplementary Table 18: SNP-based heritability of simulated traits from pooled dataset with correlated maternal-fetal genetic effects (average correlation = -0.5)

$\hat{h}^2$ of simulated traits from pooled dataset with correlated maternal-fetal genetic effects (average correlation = -0.5), estimated through conventional GCTA, M-GCTA and H-GCTA approach. Each approach was fitted using GREML (α = -0.25, -1.0), LDAK-Thin (α = -0.25, -1.0) and LDAK-Weights (α = -0.25, -1.0). For GCTA, M is the GRM generated from maternal genotypes (m), and F is the GRM generated from fetal genotypes (f). For M-GCTA, M’ represents the genetic relationship matrix of mothers; G represents genetic relationship matrix of children and D represents mother-child covariance matrix. For H-GCTA, M1 is the GRM generated from maternal transmitted alleles (m1), M2 is the GRM generated from maternal non-transmitted alleles (m2), and P1 is the GRM generated from paternal transmitted alleles (p1). A total of 100 replicates of each phenotype were simulated using empirical genotypes of Pooled dataset. P-values were calculated using z test statistics (two sided).

##### xix) Supplementary Table 19: SNP-based heritability of simulated traits from pooled dataset with correlated maternal-fetal genetic effects (average correlation = 1.0)

$\hat{h}^2$ of simulated traits from pooled dataset with correlated maternal-fetal genetic effects (average correlation = 1.0), estimated through conventional GCTA, M-GCTA and H-GCTA approach. Each approach was fitted using GREML (α = -0.25, -1.0), LDAK-Thin (α = -0.25, -1.0) and LDAK-Weights (α = -0.25, -1.0). For GCTA, M is the GRM generated from maternal genotypes (m), and F is the GRM generated from fetal genotypes (f). For M-GCTA, M' represents the genetic relationship matrix of mothers; G represents genetic relationship matrix of children and D represents mother-child covariance matrix. For H-GCTA, M1 is the GRM generated from maternal transmitted alleles (m1), M2 is the GRM generated from maternal non-transmitted alleles (m2), and P1 is the GRM generated from paternal transmitted alleles (p1). A total of 100 replicates of each phenotype were simulated using empirical genotypes of Pooled dataset. P-values were calculated using z test statistics (two sided).

##### xx) Supplementary Table 20: SNP-based heritability of simulated traits from pooled dataset with correlated maternal-fetal genetic effects (average correlation = 0.5)

$\hat{h}^2$ of simulated traits from pooled dataset with correlated maternal-fetal genetic effects (average correlation = 0.5), estimated through conventional GCTA, M-GCTA and H-GCTA approach. Each approach was fitted using GREML (α = -0.25, -1.0), LDAK-Thin (α = -0.25, -1.0) and LDAK-Weights (α = -0.25, -1.0). For GCTA, M is the GRM generated from maternal genotypes (m), and F is the GRM generated from fetal genotypes (f). For M-GCTA, M' represents the genetic relationship matrix of mothers; G represents genetic relationship matrix of children and D represents mother-child covariance matrix. For H-GCTA, M1 is the GRM generated from maternal transmitted alleles (m1), M2 is the GRM generated from maternal non-transmitted alleles (m2), and P1 is the GRM generated from paternal transmitted alleles (p1). A total of 100 replicates of each phenotype were simulated using empirical genotypes of Pooled dataset. P-values were calculated using z test statistics (two sided).

##### xxi) Supplementary Table 21: SNP-based heritability of simulated fetal traits with parent-of-origin effects (POEs) from pooled dataset

$\hat{h}^2$ of simulated fetal traits with POEs from pooled dataset, estimated through conventional GCTA, M-GCTA and H-GCTA approach using GREML (α = -1.0) model – a) 50% causal variants with POEs; b) 25% causal variants with POEs. POEs were incorporated by reducing the effect of m1 as compared to p1 by multiplying effects of m1 with (1 – I) where I is the imprinting factor such as 0.25, 0.50, 0.75 and 1.0. In each scenario, m1 shows either no imprinting i.e. I = 0.0 ($\mathrm{m1/p1} = 1.0/1.0$) or partial imprinting i.e. I = 0.25-0.75 ($\mathrm{m1/p1} = 0.75/1.0 - 0.25/1.0$) or complete imprinting i.e. I = 1.0 ($m1/p1 = 0.0/1.0$). For GCTA, M is the GRM generated from maternal genotypes (m), and F is the GRM generated from fetal genotypes (f). For M-GCTA, M' represents the genetic relationship matrix of mothers; G represents genetic relationship matrix of children and D represents mother-child covariance matrix. For H-GCTA, M1 is the GRM generated from maternal transmitted alleles (m1), M2 is the GRM generated from maternal non-transmitted alleles (m2), and P1 is the GRM generated from paternal transmitted alleles (p1). A total of 100 replicates of each phenotype were simulated using empirical genotypes of Pooled dataset. P-values were calculated using z test statistics (two sided).

##### xxii) Supplementary Table 22: SNP-based heritability of gestational duration and fetal size measurements at birth using all polymorphic SNPs

Comparison of $\hat{h}^2$ estimated through conventional GCTA, M-GCTA and H-GCTA approach for a) gestational duration, b) birth weight, c) birth length and d) head circumference. Each approach was fitted using GREML (α = -0.25), LDAK-Thin (α = -1.0) and LDAK-Weights (α = -1.0). For GCTA, M is the GRM generated from maternal genotypes (m), and F is the GRM generated from fetal genotypes (f). For M-GCTA, M' represents the genetic relationship matrix of mothers; G represents genetic relationship matrix of children and D represents mother-child covariance matrix. For H-GCTA, M1 is the GRM generated from maternal transmitted alleles (m1), M2 is the GRM generated from maternal non-transmitted alleles (m2), and P1 is the GRM generated from paternal transmitted alleles (p1). Gestational duration was adjusted for fetal sex and fetal size measurements at birth were additionally adjusted for gestational duration up to third orthogonal polynomial. Analyses using GCTA and M-GCTA approach were adjusted for 20 PCs and H-GCTA approach was adjusted for 30 PCs (10 PCs corresponding to m1, m2 and p1 each). P-values were calculated using z test statistics (one sided).

##### xxiii) Supplementary Table 23: SNP-based heritability of gestational duration and fetal size measurements at birth using SNPs with MAF > 0.001

Comparison of $\hat{h}^2$ estimated through conventional GCTA, M-GCTA and H-GCTA approach for a) gestational duration, b) birth weight, c) birth length and d) head circumference. Each approach was fitted using GREML (α = -0.25, -1.0), LDAK-Thin (α = -0.25, -1.0) and LDAK-Weights (α = -0.25, -1.0). For GCTA, M is the GRM generated from maternal genotypes (m), and F is the GRM generated from fetal genotypes (f). For M-GCTA, M' represents the genetic relationship matrix of mothers; G represents genetic relationship matrix of children and D represents mother-child covariance matrix. For H-GCTA, M1 is the GRM generated from maternal transmitted alleles (m1), M2 is the GRM generated from maternal non-transmitted alleles (m2), and P1 is the GRM generated from paternal transmitted alleles (p1). Gestational duration was adjusted for fetal sex and fetal size measurements at birth were additionally adjusted for gestational duration up to third orthogonal polynomial. Analyses using GCTA and M-GCTA approach were adjusted for 20 PCs and H-GCTA approach was adjusted for 30 PCs (10 PCs corresponding to m1, m2 and p1 each). P-values were calculated using z test statistics (one sided).

#### xxiv) Supplementary Table 24: SNP-based heritability of gestational duration and fetal size measurements at birth using SNPs with MAF > 0.01

Comparison of $\hat{h}^2$ estimated through conventional GCTA, M-GCTA and H-GCTA approach for a) gestational duration, b) birth weight, c) birth length and d) head circumference. Each approach was fitted using GREML (α = -0.25, -1.0), LDAK-Thin (α = -0.25, -1.0) and LDAK-Weights (α = -0.25, -1.0). For GCTA, M is the GRM generated from maternal genotypes (m), and F is the GRM generated from fetal genotypes (f). For M-GCTA, M' represents the genetic relationship matrix of mothers; G represents genetic relationship matrix of children and D represents mother-child covariance matrix. For H-GCTA, M1 is the GRM generated from maternal transmitted alleles (m1), M2 is the GRM generated from maternal non-transmitted alleles (m2), and P1 is the GRM generated from paternal transmitted alleles (p1). Gestational duration was adjusted for fetal sex and fetal size measurements at birth were additionally adjusted for gestational duration up to third orthogonal polynomial. Analyses using GCTA and M-GCTA approach were adjusted for 20 PCs and H-GCTA approach was adjusted for 30 PCs (10 PCs corresponding to m1, m2 and p1 each). P-values were calculated using z test statistics (one sided).

#### xxv) Supplementary Table 25: Replication of heritability estimation of gestational duration

$\hat{h}^2$ of gestational duration in HARVEST dataset based on SNPs with MAF > 0.01estimated through H-GCTA using GREML (α = -1.0) model. Gestational duration was adjusted for fetal sex. P-values were calculated using z test statistics (one sided).
